# Site-specific steric control of SARS-CoV-2 spike glycosylation

**DOI:** 10.1101/2021.03.08.433764

**Authors:** Joel D. Allen, Himanshi Chawla, Firdaus Samsudin, Lorena Zuzic, Aishwary Tukaram Shivgan, Yasunori Watanabe, Wan-ting He, Sean Callaghan, Ge Song, Peter Yong, Philip J. M. Brouwer, Yutong Song, Yongfei Cai, Helen M. E. Duyvesteyn, Tomas Malinauskas, Joeri Kint, Paco Pino, Maria J. Wurm, Martin Frank, Bing Chen, David I. Stuart, Rogier W. Sanders, Raiees Andrabi, Dennis R. Burton, Sai Li, Peter J. Bond, Max Crispin

## Abstract

A central tenet in the design of vaccines is the display of native-like antigens in the elicitation of protective immunity. The abundance of N-linked glycans across the SARS-CoV-2 spike protein is a potential source of heterogeneity between the many different vaccine candidates under investigation. Here, we investigate the glycosylation of recombinant SARS-CoV-2 spike proteins from five different laboratories and compare them against infectious virus S protein. We find patterns which are conserved across all samples and this can be associated with site-specific stalling of glycan maturation which act as a highly sensitive reporter of protein structure. Molecular dynamics (MD) simulations of a fully glycosylated spike supports a model of steric restrictions that shape enzymatic processing of the glycans. These results suggest that recombinant spike-based SARS-CoV-2 immunogen glycosylation reproducibly recapitulates signatures of viral glycosylation.

## Introduction

The Coronavirus Disease 2019 (COVID-19) pandemic has prompted the development of an unprecedented array of vaccine candidates against the causative pathogen, Severe acute respiratory syndrome-coronavirus-2 (SARS-CoV-2). All approaches aim to deliver molecular features of the virus in order to induce immunity. The viral spike glycoprotein, also referred to as S protein, has emerged as the principal focus of vaccine design efforts as antibodies against this target can offer robust immunity^1–6^. Encouragingly, neutralization can readily occur despite the extensive array of N-linked glycans distributed across the viral spike consistent with numerous vulnerabilities in this so-called glycan shield^7^. Despite these observations, glycosylation has emerged as an important parameter in vaccine development for SARS-CoV-1 and SARS-CoV-2^8,9^. The glycosylation processing state can influence immunogen trafficking in the lymphatic system^10^, influence the presentation of both native and unwanted cryptic epitopes^11^, and reveal to what extent immunogens recapitulate native viral architecture. Evidence is also emerging that glycosylation can somewhat influence the interaction between SARS-CoV-2 and its target receptor, angiotensin converting enzyme 2 (ACE2)^12–15^. The SARS-CoV-2 S protein contains at least sixty-six N-linked glycosylation sequons that direct the attachment of host glycans to specific Asn residues. This extensive glycosylation is important in lectin-mediated protein folding and direct stabilization of the protein fold^16^. In addition, certain glycans are incompletely matured during biogenesis and can lead to the presentation of immature glycans terminating with mannose residues that can act as ligands for innate immune recognition^17–19^.

Despite the focus on the S protein in vaccine development efforts, there has been considerable divergence in the mechanisms of delivery. In one approach, nucleic acid encoding the spike is delivered through mRNA or with a viral-vector^4–6,20–24^. The resulting S protein is assembled and glycosylated by host tissue. In a contrasting approach, the S protein can be recombinantly manufactured either as recombinant protein using mammalian or insect cell lines or using inactivated virus-based approaches which allows detailed characterization of the immunogen prior to delivery ^25–29^. Immunogen glycosylation can be influenced by factors specific to the manufacturing conditions such as cell type or cell culture conditions^30,31^, however, construct design and protein architecture can also have substantial impact. For example, under-processed oligomannose-sites can occur at sites sterically hidden from the host mannosidase by the tertiary or quaternary architecture including obfuscation by neighboring protein and glycan structure^18,32^. Immunogens displaying native-like architecture recapitulate these sites of oligomannose glycosylation. Conversely, immunogen design can adversely impact the presentation of native-like glycosylation. Importantly, despite the differences in biosynthesis of S protein in virions and from mammalian expression systems they seem to generate broadly similar glycosylation^33^. However, the success of a broad range of different vaccine platforms exhibiting different S protein glycosylation, indicate that native-like glycosylation is not a prerequisite for a successful vaccine. Despite this observation, understanding S protein glycosylation will help benchmark material employed in different serological and vaccine studies and help define the impact of this extensive feature of the protein surface.

The flexible and heterogeneous nature of N-glycosylation necessitates auxiliary methodologies in addition to cryo-electron microscopy or X-ray crystallography to characterize this key part of the S protein structure. Site-specific glycan analysis employing liquid chromatography-mass spectrometry is a widely used approach to obtain this information^34–38^. As research into the structure and function of the SARS-CoV-2 S protein has progressed, more details of the glycan shield of S protein have become apparent. Analyses on recombinant trimeric S protein revealed divergent N-linked glycosylation from host glycoproteins with the presence of under-processed oligomannose-type glycans at several sites^7,15,39^. Comparative analyses with monomeric and trimeric S proteins have revealed site-specific differences in glycosylation with regards to both oligomannose-type glycans and the presentation of sialic acid^39^. Analysis of S protein from insect cells demonstrated that oligomannose-type glycans were conserved on trimeric S protein, notably at N234 ^40^. In addition, molecular dynamics (MD) studies have proposed that the N234 site plays a role in stabilizing the receptor binding domain (RBD) in an exposed “up” conformation ^32^. The presence of these under-processed oligomannose-type glycans on both mammalian and insect-derived S protein provides an indication that the structure of the S protein is driving the presentation of these glycans. Subsequent studies have investigated the presentation of N-linked glycans on S protein produced for vaccination, notably the Novavax full length S protein and S protein isolated following administration of the ChAdOx-nCoV-19 vector^40,41^. The observed glycan signatures were broadly in agreement with previous analyses. However, these studies involved the truncation of glycan structures using glycosidase treatment – which is useful for categorizing glycans into high mannose or complex-type glycans and determining potential N-glycosylation sites (PNGS) occupancy on low amounts of material – but does not allow for the identification of changes in terminal glycan processing such as sialylation. The glycan processing of the two N-linked glycan sites located on the RBD has also been investigated for monomeric RBD. These sites present high levels of complex-type glycans^42,43^ and as the majority of antibodies raised against SARS-CoV-2 S protein target the RBD it is important to fully characterize the structure of the RBD, including the presentation of glycans.

N-linked glycans are highly dynamic and can substantially vary in chemical composition within a single batch of protein. Whilst glycosylation is heterogeneous, it is important for therapeutic and vaccine design to understand whether similar glycoforms arise across different protein expression platforms from different sources to ensure the antigenic surface of the S protein remains consistent when used as an immunogen or in serological assays ^44–47^.This is particularly important as glycan processing can be impacted by adverse protein conformations ^48^. Here, we describe an integrated approach including liquid chromatography-mass spectrometry (LC-MS) and MD simulations to understand the glycosylation features across recombinant S proteins from different sources. We then explore the extent to which these recombinant proteins reproduce the glycosylation of viral derived SARS-CoV-2 S protein produced from cultured Vero cells which was obtained from a previous study ^33^. We have employed an identical analytical approach to determine the glycan composition at each site, illustrating conservation across recombinant protein and virion derived material. Furthermore, we have analyzed the glycosylation of monomeric RBD recombinant protein, as it has previously been explored as a subunit vaccine and a candidate for serological testing^49,50^. We then compared the site-specific glycan analysis of the two sites located in the RBD of SARS-CoV-2, N331 and N343, between monomer and trimer and reveal a broad consensus in glycan processing, with some modest change in the processing of complex-type glycans. We also performed a comparative analysis on MERS-CoV RBD. This contrasts with glycan processing of SARS-CoV-2 RBD monomer when compared with S protein, with trimeric MERS-CoV RBD glycan sites presenting restricted glycan processing, likely due to conformational masking of these sites, either by proximal glycans or nearby protein clashes, on trimeric MERS-CoV S protein ^51,52^. We have further combined the site-specific glycan data of SARS-CoV-2 S protein with MD simulations. These simulations reveal distinct degrees of accessibility between different glycan sites across the protein that broadly correlate to their processing states. Taken together, our results reveal the conserved structural N-glycan sites in S protein when compared with native viral spike which drives similarity in glycosylation amongst S protein from disparate sources and manufacture methods. Understanding S protein glycosylation will aid in the analysis of the vaccine and serological work of the global COVID-19 reponse.

## Results

### Expression and purification of recombinant S protein from multiple sources

In order to define the variability in trimeric recombinant S protein glycosylation and compare recombinant and viral derived S protein we obtained preparations of recombinant S protein from a range of laboratories. These include the Amsterdam Medical Centre (Amsterdam), Harvard Medical School (Harvard), Switzerland, The Wellcome centre for Human genetics (Oxford) and Biological Sciences/University of Texas at Austin (Southampton/Texas)^1,29,53–56^. Whilst there are minor differences between the constructs used to produce the S protein the overall design is similar. This involves a truncation of the S protein prior to the C terminus, replacement of the furin cleavage site between S1 and S2 with a “GSAS” or otherwise mutated linker, a C-terminal T4 fibritin trimerization motif and stabilization of the prefusion S protein conformations using proline substitutions^29,51,57^. The recombinant proteins analyzed contain 22 N-linked glycan sites with the exception of the “Amsterdam” preparation which was truncated prior to the last 3 N-glycans. An alignment of the recombinant proteins was performed and glycan sites are numbered according to the numbering used in previous reports of the site-specific glycosylation of the 2P stabilized S protein (**Supplementary Figure 1**) ^7^. All recombinant proteins were expressed in human embryonic kidney 293 (HEK293) cells with the Amsterdam, Harvard, and Southampton/Texas S protein preparations produced in HEK293F and the Oxford S protein preparion in HEK293T. The “Swiss” S protein was expressed in Chinese hamster ovary (CHO) cells. Following expression and purification, an identical approach was utilized to prepare the recombinant samples for analysis by LC-MS, involving three separate protease digests; trypsin, chymotrypsin and alpha lytic protease. After analysis by LC-MS the site-specific glycosylation of these recombinant samples was compared to previously published viral derived S protein which we searched using the same analytical parameters as for the recombinant proteins ^33^.

### Conservation of under-processed glycans on trimeric recombinant and viral-derived S protein

Under-processed oligomannose-type and hybrid-type glycans are common on viral glycoproteins and commonly arise due to either glycan- or protein-mediated steric clashes with ER and Golgi resident mannosidase enzymes, terminating the glycan processing pathway. As the presence of these glycans is linked to the quaternary structure of the protein, changes in the abundance of these glycoforms can indicate changes in the fine structure of the glycoprotein. To investigate the abundance of these glycans, we simplified the heterogeneous glycan compositions detected by LC-MS into three categories, as (1) oligomannose-type and hybrid-type glycans, (2) complex-type glycans and (3) the proportion of PNGS lacking an N-linked glycan. Overall, the recombinant samples recapitulated the glycan processing observed on the viral derived S protein (**Figure 1A**). Sites on the viral-derived material with an abundance of under-processed glycans greater than 30% include N61, N122, N234, N603, N709, N717, N801 and N1074 (**Figure 1B**). With a few exceptions, all recombinant samples analyzed also present at least 30% oligomannose-type glycans at these sites (**Supplementary Table 1**).

**Figure 1:**
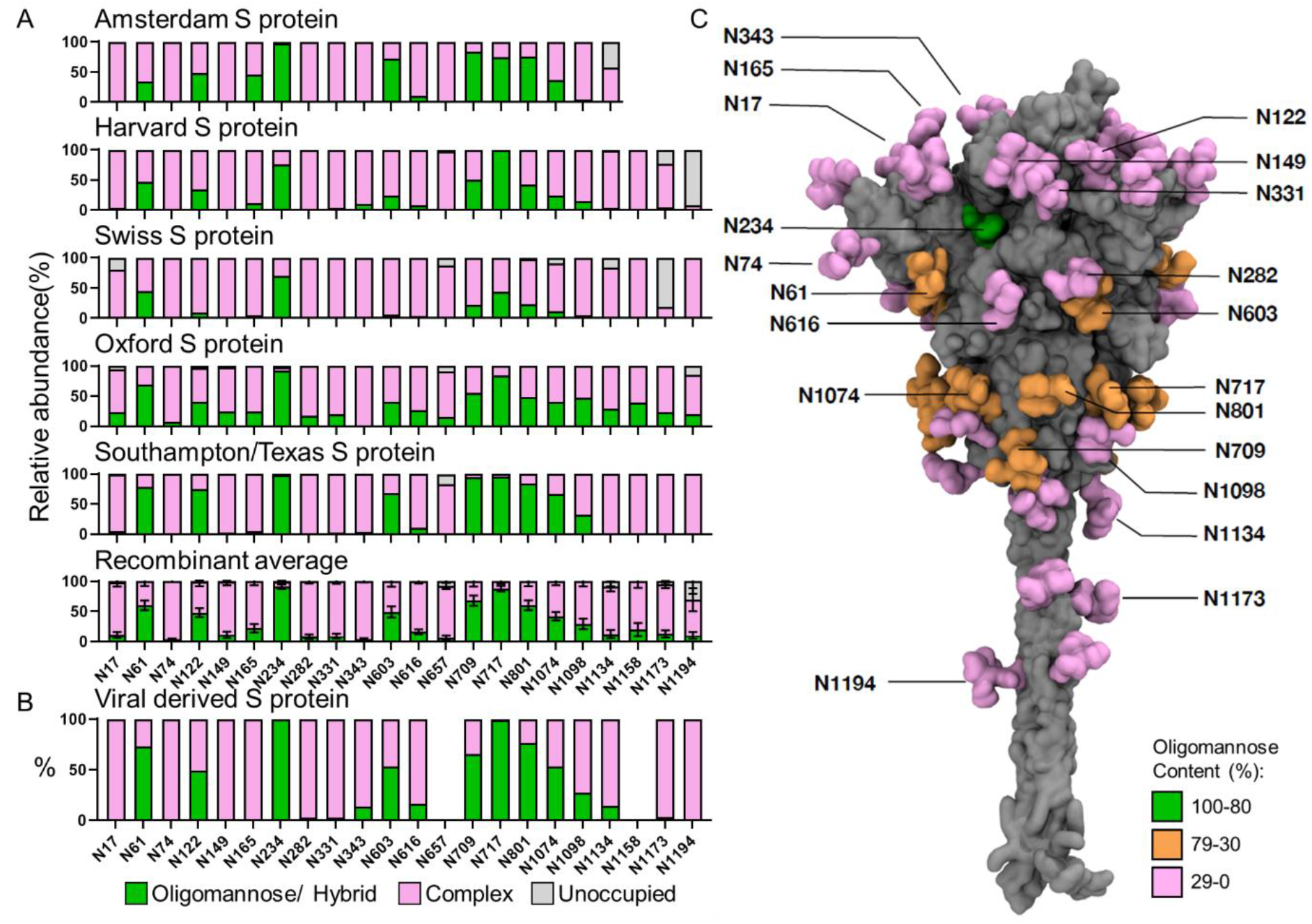
The site-specific glycosylation of recombinant and viral derived S protein from multiple laboratories. **(A)** Site-specific glycan analysis of recombinant S protein expressed and purified at different locations. The bar charts represent the relative proportion of glycoforms present at each site, including the proportion of PNGS that were not modified by an N-linked glycan. The proportion of oligomannose- and hybrid-type glycans are colored green. Processed complex-type glycans are colored pink and the proportion of unoccupied sites are colored grey. The institution which provided the S protein for analysis is listed above each chart. The Texas/Southampton data is reproduced from Chawla et al. (unpublished data). The average compositions of recombinant S protein were calculated using all samples. Bars represent the mean +/- standard error of the mean of all recombinant samples analyzed. **(B)** The viral derived site-specific analysis was obtained from data acquired by Yao et al. and categorized in the same manner as above^33^. Data for sites N657 and N1158 could not be obtained and are not represented. **(C)** Full length model displaying the site-specific oligomannose glycosylation of viral derived S protein; description of how this model was generated can be found in **Materials and Methods**. Both protein and glycans are shown in surface representation; the former coloured in grey and the latter coloured based on oligomannose content as shown in **Supplementary Table 1** (green for 80-100%, orange for 30-80% and pink for 0-30%).

To compare the variability of oligomannose- and hybrid-type glycosylation across all recombinant proteins the compositions were averaged and displayed with the standard error of the mean (SEM) of S protein preparations from different laboratories (**Figure 1A** and **B**). This analysis revealed a broad consensus of glycan processing regarding high mannose glycans, with localized variations occurring. One example of remarkable homogeneity is the N234 site, which presents high levels of under-processed glycans in all samples analyzed. This processing was conserved on the viral derived S protein. The recombinant material possessed low levels of oligomannose-type glycans at several sites across the protein not present on viral derived material. For example, material analyzed from Oxford had at least 5% oligomannose-type glycans at every site except N343 (**Figure 1A**). This global moderate increase in oligomannose-type glycans could potentially arise from two sources, the first is that the recombinant preparations utilize stabilizing mutations that might generate a more compact structure which limits the ability of glycan processing enzymes to act, this is supported by the observation that viral-derived S protein exhibits metastable conformations in the NTD, likely increasing accessibility to particular glycan sites ^33^. The second could be that the higher yield of protein obtained during recombinant protein expression places limitations on the capacity of the cell to process the high number of glycan sites present on the S protein.

Glycan occupancy is another important parameter to consider for immunogen design. Underoccupancy of recombinant glycoprotein PNGS compared to their viral counterparts has been shown to result in the presentation of non-neutralizing distracting epitopes and is important to monitor^58,59^. As is the case with viral derived HIV-1 Env, viral derived SARS-CoV-2 S protein displayed high levels of glycan occupancy at all sites analyzed (**Figure 1B**). When comparing this to the consensus site-specific glycosylation data, the majority of PNGS on recombinant S protein are highly occupied. The exceptions to this are at the C terminus, where several samples analyzed display reduced glycan occupancy denoted by a larger population of unoccupied sites (**Figure 1A**). These sites displaying larger proportions of unoccupied glycans at the C terminus is likely due to the truncation of the recombinant proteins required for solubilization; it has previously been reported that the proximity of a glycan sequon to the C-terminus can influence its occupancy^60^.

Modelling the site-specific glycosylation of viral derived material enables the glycosylation of S protein to be contextualized spatially in three dimensions. Using the cryo-EM structure of S protein ectodomain (ECD) in the open state (one RBD in the “up conformation” and two RBDs in the “down conformation”) (PDB: 6VSB)^29^, as well as the NMR structures of SARS-CoV HR2 domain (PDB: 2FXP)^61^ and HIV-1 gp-41 TM domain (PDB: 5JYN)^62^, a complete model of the S protein was generated (details in Methods section). This model includes glycosylation sites which are often not resolved as they are present on variable loops or along the flexible stem region of the S protein. This model demonstrated the spread of different glycan processing states across the S protein (**Figure 1C**). As with other published models of spike glycosylation, the N234 site is extensively buried within the protein surface proximal to the RBD and rationalizes the conserved presentation of oligomannose-type glycans on recombinant and viral-derived material as the glycan processing enzymes are unable to completely process these sites. Glycan sites located on variable loops and on the exposed stem of the S protein are much more processed as these sites are more exposed.

### Divergent glycan processing at sites presenting complex-type glycans

Whilst glycan under-processing can provide information pertaining to the quaternary structure of the glycoprotein, the majority of glycans across the S protein are able to be processed beyond these glycoforms. Glycan sites not under the same structural pressures as at the restricted sites such as N234 will be processed in an analogous manner as those presented on the majority of host glycoproteins. By comparing the site-specific glycan compositions at example sites, we sought to understand the variability in glycan elaboration across multiple preparations of S protein. For this analysis we compared the recombinant protein glycosylation data presented in this study (Amsterdam, Harvard, Swiss and Oxford) with viral-derived S protein previously reported, but analyzed using the same search parameters as for the recombinant proteins. To aid comparison, we selected one site which presented high levels of under-processed oligomannose-type glycans (N234), one that presented a mixture of processing states (N1074), and a site that was populated by complex-type glycans (N282) (**Figure 2**). Comparing the site-specific glycosylation of N234 demonstrated the conservation of oligomannose-type glycan processing. On 4 of the 5 samples presented in **Figure 2** the most abundant glycan detected was Man8GlcNAc2, and on the Amsterdam S protein Man9GlcNAc2 predominated, which is marginally less processed. The Harvard and Swiss samples displayed elevated glycan processing at N234, but these glycans were only ∼20% abundant and were still the least processed site on these samples (**Figure 2** and **Supplementary Table 1**).

**Figure 2:**
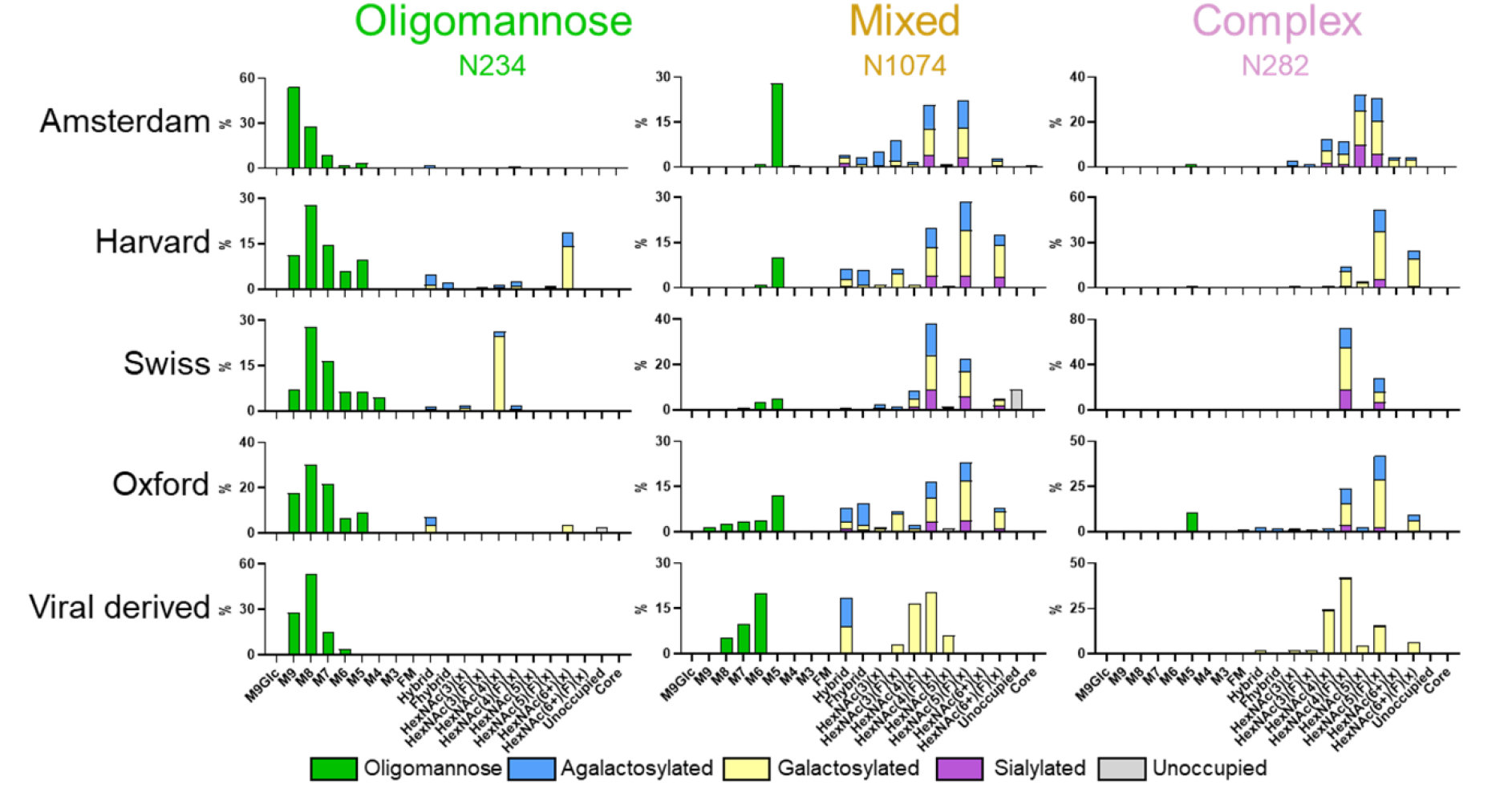
Detailed composition comparison between sites with differential processing states. Glycan compositions at N234, N1074 and N282. For all samples analyzed, glycans were categorized and colored according to the detected compositions. Oligomannose-type glycans (M9-M4) were colored green. Hybrid-type glycans, those containing 3 HexNAc and at least 5 hexoses, were colored as for complex-type glycans since one arm can be processed in a similar manner. Complex-type glycans were categorized according to the number of HexNAc residues detected and the presence or absence of fucose. Core glycans represent any detected composition smaller than HexNAc2Hex3. For hybrid and complex-type glycans bars are colored to represent the terminal processing present. Blue represents agalactosylated, yellow is galactosylated (containing at least one galactose) and purple is sialylated (containing at least one sialic acid). The proportion of unoccupied PNGS is colored grey.

The N1074 site presented a diverse mixture of oligomannose-, hybrid-, and complex-type glycans across all samples analyzed. The abundance of oligomannose-type glycans was more variable at this site with the total oligomannose-type glycan content varying from 11% to 35% (**Figure 2**). The predominant composition at N1074 varied from sample to sample: for the Amsterdam recombinant preparation and the viral derived S protein, an oligomannose-type glycan was the most abundant glycan at N1074; however, for the other samples a fully processed glycan was more abundant. For all samples analyzed, N1074 presented diverse glycan processing states. The elaboration of complex-type glycans with different monosaccharides can influence the function of the glycoprotein to which they are attached, for example, sialylation can extend the half-life of a glycoprotein in the body^63^. As they progress through the Golgi, glycans can be elaborated by the addition of fucose, galactose and sialic acid. The abundance of these monosaccharides is influenced by the cell from which they are produced and by the culture conditions or media compositions under which recombinant proteins are manufactured. Typically, glycoproteins produced from HEK293F and CHO cells present complex-type glycans with high levels of fucosylation and galactosylation but lower levels of sialylation ^64,65^. The processing of N1074 and N282 demonstrates this with the majority of complex-type glycans bearing at least one fucose on all recombinant S preparations. Likewise, the majority of glycans are galactosylated, although populations of glycans are present which lack any elaboration beyond N-acetylglucosamine branching. Hybrid-type glycans, where one arm of the glycan is processed and elaborated as for complex-type glycans and one remains under-processed, are present in lower abundances across N1074 in recombinant preparations. These hybrid-type glycans are also generally of low abundance on mammalian glycoproteins ^66^.

Whilst the complex- and hybrid-type glycan compositions are variable, there are some visible trends when comparing glycan processing between recombinant and viral-derived S protein. The starkest difference is the lack of sialic acid residues across not only N1074 and N282 but across all PNGS of viral derived material (**Figure 2, Supplementary Table 1**). Upon averaging all the recombinantly produced S protein and comparing to viral derived S protein there is a 21 percentage-point decrease in sialylation (**Supplementary Table 2**). The fucosylation of viral derived material is also lower across both N1074 and N282 and this trend is again mirrored across all sites, with viral derived S protein possessing a 16 percentage-point decrease in fucosylation (**Supplementary Table 2**). The final difference is in glycan branching. The number of processed glycan antenna can be inferred from the detected number of N-Acetylhexosamine (HexNAc) residues determined by LC-MS. The more HexNAc present on a glycan composition the more branched that glycan tends to be, for example HexNAc(6) can correspond to a tetra-antennary glycan whereas HexNAc(4) is biantennary. As glycan processing is heterogeneous, this change is more subtle, however the complex-type glycans of N1074 and N282 are less branched on viral S protein compared to the recombinant S proteins, with a ∼30 percentage point decrease in abundance of HexNac(5) and HexNAc(6) corresponding to tri- and tetra-antennary glycans (**Figure 2**). These changes are also apparent on other processed sites such as N165 and N1158 (**Supplementary Table 2**).

These changes contrast expected observations based upon similar analyses comparing viral derived Env and soluble recombinant variants. For HIV-1 Env, an increase in glycan branching and sialylation is observed across sites presenting complex-type glycans^67^. These changes were also present on recombinant full length Env^68^. Several factors could be influencing these changes. The first is that the membrane tether afforded to the viral Env brings the glycans into close proximity to the glycan processing enzymes and the second is that the producer cell had greater expression levels of the glycosyltransferase enzymes involved in glycan processing. The SARS-CoV-2 S protein in a viral context is likely under similar constraints; however, the glycan processing is distinct. One factor which could be important is the early budding of the SARS-CoV-2 virion into the ER/Golgi intermediate complex (ERGIC) which may distance the spike protein from glycosyltransferases present in the trans-Golgi compared to Env, which remains attached to the membrane, proximal to glycosyltransferases^69^. The choice of producer cell and culture condition of the virus, in this case Vero cells, which are derived from the kidney cells of *Cercopithecus aethiops* (African green monkey), may also influence the glycan processing as the expression levels of glycosyltransferase enzymes may account for the diminished attachment of sialic acid and fucose observed on viral S protein compared to recombinant S protein. As these changes in glycosylation likely are not impacting the immunogenicity of the viral spike mimetics, as evidenced by the high efficacy of several vaccine candidates, these observations still remain important when considering how the virus may be interacting with the immune system via lectin interactions, and may be informative when considering antigenic tests and purifications using glycan binding reagents.

### The expression of monomeric RBD constructs impacts glycan processing

Our analysis demonstrates that S protein glycosylation is influenced by quaternary protein architecture, and other factors. We next sought to compare the glycosylation of soluble recombinant monomeric RBD with that of recombinant and viral derived trimeric S protein. We expressed and purified recombinant RBD and compared the site-specific glycosylation of the two glycan sites located in the RBD, N331 and N343, with those observed on recombinant S protein and viral derived S protein reported previously^7,33^. Overall, the N331 and N343 glycans across all expression formats were highly processed, with little to no oligomannose-type glycans detected (**Figure 3A** and **B;** and **Supplementary Table 3**). For the RBD sites presented on viral derived S protein, the complex-type glycans observed were similar to those observed at N282 with most of the PNGS occupied by bi- and tri-antennary glycans (HexNAc(4) and HexNAc(5)) (**Figure 3A**). The majority of these glycans were fucosylated, however, a large proportion of the complex type-glycans on viral derived S lacked fucosylation (24% N331 and 20% N343). As with other sites on the viral derived S protein, minimal levels of sialylation were observed and the majority of glycans possessed at least 1 galactose residue.

**Figure 3:**
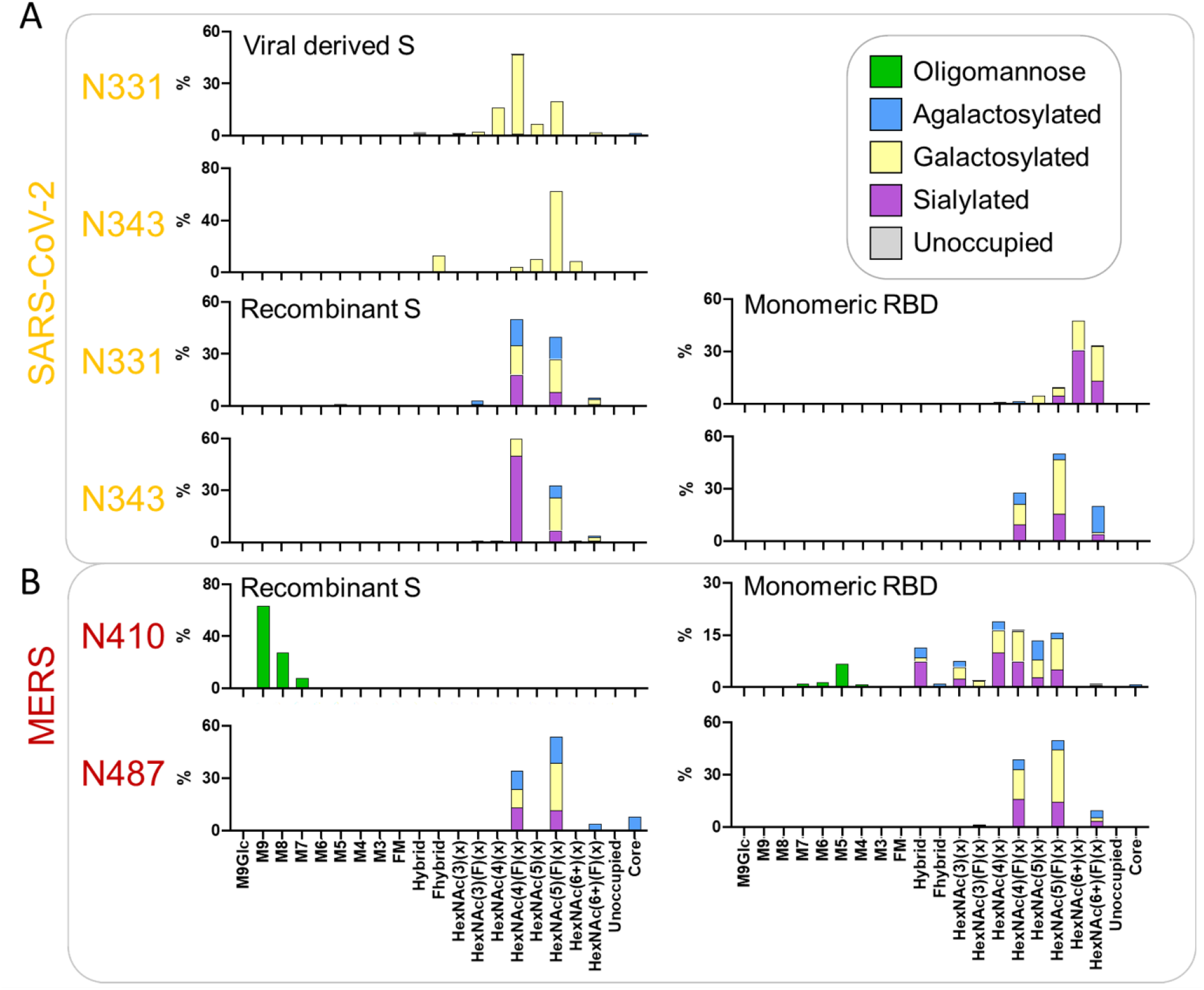
Comparative analysis of the glycosylation of the two PNGS on the RBD for viral derived S protein, recombinant S protein and monomeric RBD. **(A)** Detailed site-specific glycan compositions of the two sites located in the RBD of SARS-CoV-2. Recombinant S protein data is reproduced from Chawla et al. (unpublished) and data for the viral S protein RBD sites was obtained from Yao et al.^33^. Site-specific glycan data is presented as outlined in Figure 2. **(B)** Site-specific compositions for N-glycan sites located in the RBD of MERS-CoV when expressed as part of a soluble recombinant S protein compared to RBD-only. Data for the MERS-CoV S protein were obtained from a previous study ^52^. Site-specific glycan data is presented as outlined in Figure 2.

In comparison, the two RBD sites of recombinant soluble S protein are highly fucosylated, with close to 100% of the glycans at N331 and N343 containing at least one fucose (**Figure 3B**). As with the viral-derived S protein the majority of the glycans are bi- and tri-antennary complex-type glycans. The recombinant S protein RBD sites are also more sialylated than N331 and N343 on viral-derived S protein, with 28% of N331 glycans containing at least one sialic acid and 60% of those on N343 (**Supplementary Table 3**). Interestingly, when RBD is expressed as a monomer there are additional subtle changes compared to viral derived and recombinant S protein. The most prominent change is in the glycan branching; whereas recombinant and viral derived RBD glycan sites possess low quantities of tetra-antennary glycans approximately, one third of the glycans at N331 and one fifth of the glycans at N343 consist of these larger branched structures (**Figure 3**). When compared to recombinant S protein the monomeric RBD sites also possess low levels of biantennary glycans. Despite these changes both recombinant trimeric S protein and monomeric RBD sites have high levels of fucosylation. These results suggest that complex-type glycans are under differential control hierarchies where certain forms of glycan processing could be influenced by the structural presentation of glycan sites. The attachment of fucose, galactose and sialic acid for these RBD sites appears to be controlled by more global phenomena, such as the producer cell, as the attachment of these monosaccharides is similar when comparing monomeric RBD and trimeric recombinant S protein which were produced in identical cell lines. The branching of the RBD sites on monomeric RBD is greater than that of trimeric S protein, both viral and recombinant, and suggests that the quaternary structure of the glycoprotein may have a small role on the elaboration of complex-type glycans of the RBD. These results are similar to previous analyses that have compared trimeric S protein with monomeric S1^39^.

To further explore differences in glycosylation of RBD PNGS, we performed a similar comparative analysis on MERS-CoV. The site-specific glycan analysis of recombinant MERS-CoV S protein has been reported previously^52^. For the comparative analysis of trimeric recombinant protein and RBD, the MS files obtained in the previous study were searched using an identical version of the analysis software using the same glycan libraries as for the analysis displayed in **Figure 1 and 2**. This analysis revealed differences at the glycan sites present on MERS RBD when expressed monomerically compared to recombinant trimeric soluble S protein. One of the sites, N487, is similar between the two platforms, presenting glycoforms typical of sites populated by complex-type glycans (**Figure 3B**). In contrast N410 is occupied by exclusively oligomannose-type glycans when present on recombinant trimeric S protein. When monomeric MERS RBD is expressed the processing of the N410 site is markedly increased, with complex-type glycans predominating the glycan profile, although a subpopulation of oligomannose-type glycans remain (**Figure 3B**). Modelling the N410 glycan onto published structures of MERS S protein reveals that the N410 glycan is protected from processing by mannosidase enzymes such as ER α mannosidase I when buried within the trimer, but when presented on monomeric RBD it is readily accessible to glycan processing (**Supplementary Figure 2**). These observations further highlight how the quaternary structure of a glycoprotein is a key determinant of the glycan processing state and demonstrates how glycan sites can provide information about protein folding and quaternary structure.

### Molecular dynamics simulations reveal the relationship between accessibility and glycan processing

Using site-specific glycan analysis, it is possible to infer the structural restraints placed on particular PNGS by the quaternary structure of the protein. MD simulations can help to understand how the protein flexibility can influence glycan processing. The recombinant proteins analysed in this study all use proline substitutions to stabilize the pre-fusion S protein conformation. Structural analyses of S protein containing these proline substitutions have shown that one of the three subunits frequently displays its RBD in an up conformation^29^. By performing simulations using models containing one RBD up, it is possible to investigate how differential RBD presentation can impact site-specific glycosylation.

To this end, we performed 200 ns MD simulations of fully glycosylated trimeric S protein embedded within an ERGIC membrane model (details in **Materials and Methods**). The Man9GlcNAc2 glycan (Man-9), which represents the primary substrate for glycan processing enzymes, was added to each PNGS to understand the effect of protein and glycan dynamics on glycan processing. Glycan accessibility to enzymes was then elucidated by calculating the accessible surface area (ASA). To ensure the correct size of probe was used for ASA calculation, we first looked at the structure of mannosidases and glycosyltransferases bound to its substrate or substrate analogue ^70,71^. By measuring the distance between the substrate binding site and the outer surface of the enzymes, we found a probe with 1.25-1.5 nm radius would best approximate the size of the enzymes (**Supplementary Figure 3**). We measured ASA values for chain A of the S protein using probes of 1.25 and 1.5 nm radii and found they both gave very similar results. Plotting accessible points of a 1.5 nm probe around individual glycans also indicated that a 1.5 nm radius would be required for the Man-9 glycan to be accessed by mannosidases and glycosyltransferases. We therefore used a 1.5 nm probe to measure ASA of each Man-9 glycan on the S protein model during the MD simulation.

Comparing the ASA of each glycan across each protomer reveals a diverse range of accessibility (**Figure 4A and B**). Certain sites such as N234, were determined to possess low ASA values (**Supplementary Figure 4A**), indicating that this site is extremely buried, correlating with observations that N234 is the least processed on all recombinant and viral derived S protein. Conversely, the most exposed site across all three chains is N74 (**Supplementary Figure 4B**). This glycan is highly processed when analyzed by LC-MS. Interestingly, previous analyses have shown that N74 possesses sulfated glycans^15^. We have included sulfated compositions in our glycan library to search the MS data and observe that N74 contains multiple sulfated glycan compositions on all samples produced in HEK293 cells (**Supplementary Table 1**). The higher accessibility of this site may explain why sulfated glycans are observed at higher abundances at this site, but not others. Some glycans show a distinctly bimodal accessibility pattern, wherein high and low ASA values were observed along the trajectory (**Supplementary Figure 5**). For example, two of the N122 glycans were buried within a crevice between the N-terminal domain of one chain and the RBD of an adjacent chain during a portion of the simulation. Similarly, one of the N603 glycans inserted into a large inter-protomeric groove near the S1/S2 cleavage site. This observation correlates with LC-MS data from both recombinant and viral derived S protein, showing that these two sites are populated by approximately half oligomannose/hybrid and half complex glycans, potentially due to the ability of these glycans to be either buried or exposed on the protein surface. The glycan on N165 also showed bimodal ASA values. However, this is due to the RBD up configuration, which allows N165 glycan to insert into the gap between the RBD and the N-terminal domain, which was not accessible in the RBD down configuration. N165, along with N234 glycans, have been shown to modulate the RBD’s conformational dynamics by maintaining the up configuration necessary for ACE2 recognition^32^.

**Figure 4:**
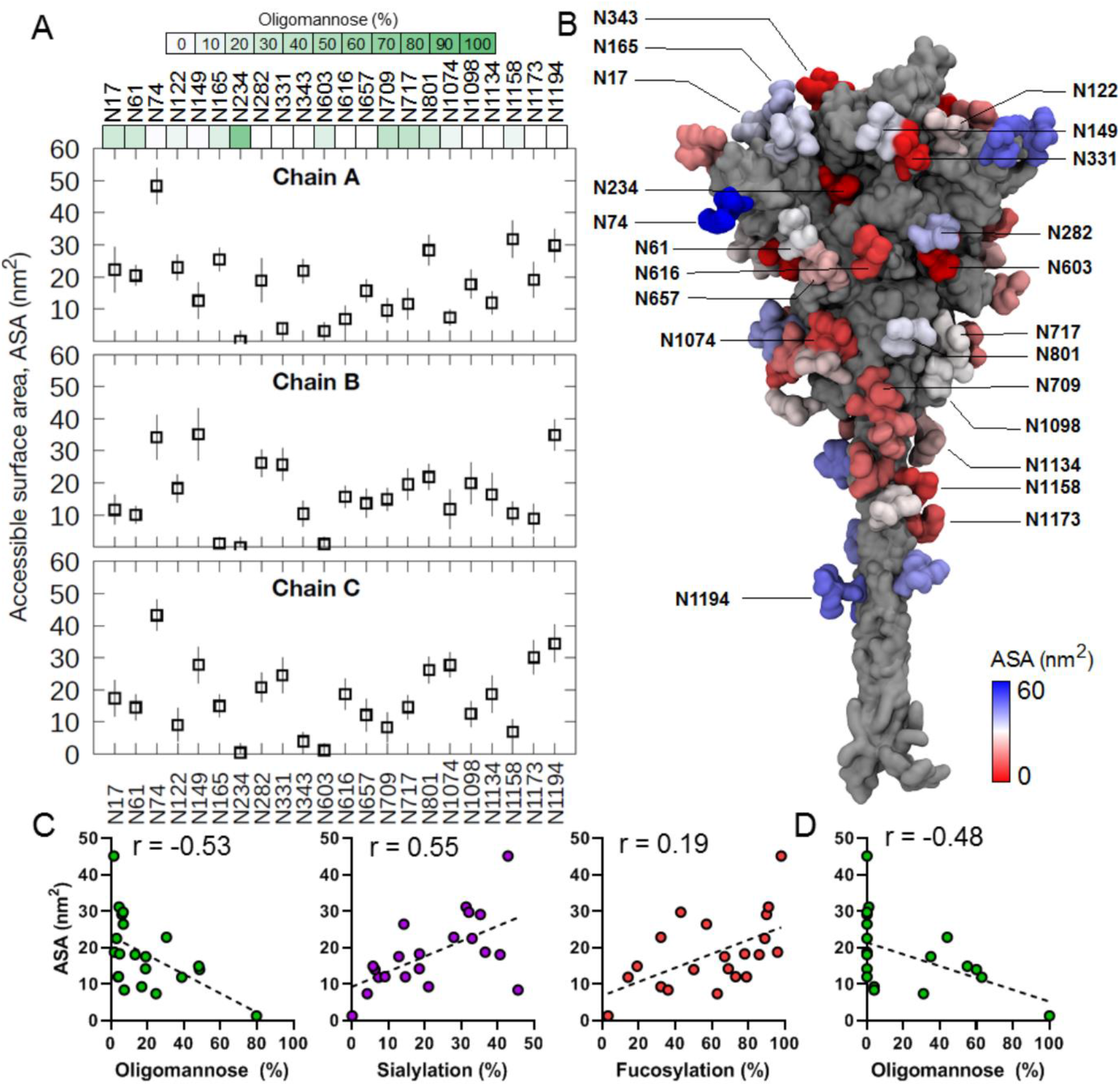
Accessible surface area (ASA) of oligomannose-type glycans from MD simulations. **(A)** The ASA values were calculated for each oligomannose-type (M9) glycan for all three S protein chains. In this model, chain A is modelled in the RBD “up” conformation. The last 50 ns of the simulation was used for calculation and error bars indicate standard deviation along the trajectory. The probe size used was 1.5 nm. The green color bars represent the oligomannose content of respective sites, calculated as the average oligomannose-type glycan content of the recombinant S proteins analyzed in this manuscript (Amsterdam, Harvard, Swiss and Oxford S proteins). **(B)** The structure of the S protein (grey) with glycans coloured based on their ASA values shown in surface representation. **(C)** Graphs comparing glycan processing to calculated ASA values, averaged for chains A, B and C, for the recombinant protein average for the oligomannose-type glycan content, proportion of glycans containing at least one sialic acid and the proportion of glycans containing at least one fucose. The reported r values represent the Spearman’s rank correlation coefficient. **(D)** The oligomannose-type glycan content versus ASA for viral derived SARS-CoV-2 S protein presented as in Panel C.

To understand the relationship between oligomannose-type glycan presentation and the accessible surface area of each glycan site we used the average oligomannose-type glycan content of each site as determined in **Figure 1**, for both the recombinant and viral derived material, and compared it to the average ASA values for chains A, B and C, as LC-MS cannot distinguish which glycan came from which chain (**Figure 4C** and **D**). For both recombinant and viral derived S protein, there was a negative correlation between ASA and oligomannose content (Spearman’s correlation coefficient -0.53 for recombinant and -0.48 for viral derived). This demonstrates the link between glycan processing and quaternary architecture, as sites which are less accessible to the glycan processing enzymes present higher populations of immaturely processed oligomannose-type glycans. Conversely, the processing of complex-type glycans such as the addition of sialic acid correlated with the accessbile surface area at a site-specific level (r = 0.55) and also fucosylation, albeit to a lesser extent (r = 0.19). Whilst there is an observed correlation between oligomannose-type glycans and ASA, the nanosecond time scale captured during MD simulations is shorter compared to the overall time taken for glycan processing in the ER and Golgi apparatus to occur, and therefore the S protein will be able to sample more conformations not resolved during simulation.

Finally, to determine the potential effect of glycan clustering on enzyme accessibility and subsequent glycan processing, we calculated the number of contacts made by each glycan at a given site with its surrounding glycans during the simulation (**Supplementary Figure 6**). Interestingly, in all three chains, the sites on the S2 subunit show a higher number of glycan-glycan contacts, suggesting that they are most likely to form local clusters. An overlay of consecutive simulation snapshots of glycans also clearly shows that the S2 region is more densely packed with glycans compared to S1 and HR2 domains. This agrees with the LC-MS data showing that the sites on the S2 subunit, specifically N709, N717 and N801, have a high degree of unprocessed oligomannose-type glycans. The high density of glycan-glycan contacts within the mannose glycan patch has previously been implicated in reduced processivity by glycan processing enzymes in HIV-1 gp120^72^, and a similar mechanism might explain increased oligomannose content at S2 sites of SARS-CoV-2. A recent study showed that the glycan-binding anti HIV-1 broadly neutralizing antibody 2G12 can bind to S protein via a common epitope around N709 on the S2 subunit^73^, further suggesting the formation of a local glycan cluster. Collectively, our MD simulation sampling shows that glycan accessibility to host enzymes, which is affected by protein quaternary structure and local glycan density, is an important determinant of glycan processing.

### Perspectives

The global impact of COVID-19 has resulted in laboratories across the globe producing recombinant spike protein for vaccine design, antigenic testing and structural characterization. Whilst the site of glycan attachment is encoded by the viral genome, the processing of the attached glycans can also be influenced by a wide range of exogenous phenomena, including recombinant host systems and processes for production in cell culture. Using immediately available materials, we have compared the glycosylations of Vero-produced virus preparations for Spike proteins with those of HEK-293 and CHO-derived recombinant Spike proteins. We are aware that these preparations are likely to be different from Spike proteins as produced in a diversity of cells in infected patients. However, here we show that when a glycan site is located in regions of the SARS-CoV-2 S protein which are not readily accessible the glycan site will possess high levels of under-processed oligomannose-type glycans, a phenomenon that will likely be of a general nature. This conclusion is derived from the observation that the under-processing of glycans is found in protein preparations from a range of institutions across the globe and are also present on Vero cell derived S protein derived from infectious SARS-CoV-2. Whilst the presence of oligomannose-type glycans are rare on the majority of host glycoproteins their abundance on SARS-CoV-2 is lower than that of other viral glycoproteins and means that the glycan shield density of this virus is low when compared to HIV-1 Env and Lassa GPC. This is likely to mean that the immunodominant protein epitopes remain exposed. The processing equivalence between recombinant and viral derived S protein indicates that recombinant S protein glycosylation for vaccinations will likely mimic those occuring in human infections and will remain antigenically comparable. By analysing complex-type glycan processing across multiple samples, we demonstrate that this glycosylation is driven more by other parameters including the producer cell and culturing conditions. One observed exception to this was that the expression of monomeric RBD as opposed to trimeric S protein does increase glycan branching on the two glycan sites located on SARS-CoV-2 RBD. This suggests that the glycan processing of complex-type glycans, in addition to oligomannose-type glycans, may also be under structural constraints, albeit to a much lesser extent. Overall, these results demonstrate that whilst N-linked glycosylation is highly diverse at certain regions of the S protein, there is a broad consensus of glycan processing with regards to oligomannose-type glycans between virus and immunogen S protein. This is something which cannot be taken for granted, as when comparing recombinant and viral derived HIV-1 Env, the reduced glycan occupancy of immunogens can induce an immune response that is incapable of neutralizing the virus. The reproducibility of S protein glycosylation from many different sources is of significant benefit for immunogen design, serology testing and drug discovery and will mean that the glycans of SARS-CoV-2 are unlikely to provide a barrier to combatting the COVID-19 pandemic.

## Acknowledgements

The authors thank Jason McLellan and Daniel Wrapp for providing material for analysis of the Southampton/Texas S protein and Florian Wurm (Prof. emeritus, EPFL, Lausanne, Switzerland) for critical reading and editing the final manuscript. This work was supported by: the International AIDS Vaccine Initiative (IAVI) through grant INV-008352/OPP1153692 and the IAVI Neutralizing Antibody Center through the Collaboration for AIDS Vaccine Discovery grant OPP1196345/INV-008813 (D.R.B and M.C.), both funded by the Bill and Melinda Gates Foundation; National Institute for Allergy and Infectious Diseases through the Scripps Consortium for HIV Vaccine Development (CHAVD) (AI144462) and the University of Southampton Coronavirus Response Fund (D.R.B. and M.C.); PJB is supported by BII of A*STAR. Simulations were performed on the petascale computer cluster ASPIRE-1 at the National Supercomputing Centre of Singapore (NSCC) and the A*STAR Computational Resource Centre (A*CRC); the National Institutes of Health grant P01 AI110657 (RWS), Bill and Melinda Gates Foundation grants OPP1132237 and INV-002022 (RWS), Vici grant from the Netherlands Organization for Scientific Research (NWO) (RWS); and the Bill and Melinda Gates Foundation (OPP 1170236 to D.R.B.); NIH grants AI147884 (B.C.), AI147884-01A1S1 (B.C), AI141002 (B.C.), AI127193 (B.C.), a COVID19 Award by Massachusetts Consortium on Pathogen Readiness (MassCPR; B.C.), as well as a Fast grant by Emergent Ventures (B.C.); and the Chinese Academy of Medical Sciences (CAMS) Innovation Fund for Medical Science (CIFMS), China (grant number: 2018-I2M-2-002) to D.I.S.; H.M.E.D. was supported by the Wellcome Trust (101122/Z/13/Z); and D.I.S. by the UK Medical Research Council (MR/N00065X/1).

## Conflict of interest

ExcellGene sells purified trimeric Spike protein preparations from CHO cells to commercial companies for internal research and for use in diagnostic applications.

## Author contributions

J.D.A, Y.W. and H.C. performed site-specific analysis, J.D.A, F.S., H.C., P.J.B., and M.C. wrote and edited the manuscript. F.S., L.Z., A.T.S., M.F. and P.J.B. performed MD simulations W.H., S.C., G.S., P.Y., R.A. and D.R.B. provided monomeric RBD proteins. P.J.M.B., Y.C., H.M.E.D, T.M., J.K., P.P., M.J.W, B.C., D.I.S, and R.W.S provided recombinant S protein. Y.S. and S.L. produced and collected LC-MS data for viral derived S protein. All authors edited and reviewed the manuscript prior to submission.

## Materials and Methods

### SARS-CoV-2 S protein purification (Harvard)

To express a stabilized ectodomain of Spike protein, a synthetic gene encoding residues 1−1208 of SARS-CoV-2 Spike with the furin cleavage site (residues 682– 685) replaced by a “GGSG” sequence, proline substitutions at residues 986 and 987, and a foldon trimerization motif followed by a C-terminal 6xHisTag was created and cloned into the mammalian expression vector pCMV-IRES-puro (Codex BioSolutions, Inc, Gaithersburg, MD). The expression construct was transiently transfected in HEK 293T cells using polyethylenimine (Polysciences, Inc, Warrington, PA). Protein was purified from cell supernatants using Ni-NTA resin (Qiagen, Germany), the eluted fractions containing S protein were pooled, concentrated, and further purified by gel filtration chromatography on a Superose 6 column (GE Healthcare).

### SARS-CoV-2 S protein purification (Oxford)

The extracellular domain of SARS-CoV-2 spike was cloned into pHLsec vector^74^ and encompassed residues M1-Q1208 (UniProtKB ID P0DTC2) with mutations R682G/R683S/R685S (furin recognition sequence), K986P/V987P, followed by Fibritin trimerisation region, an HRV3C protease cleavage site, a His8-tag and a Twin-Strep-tag at the C-terminus^29,55^. Spike ectodomain was expressed by transient transfection HEK293T (ATCC, CRL-3216) cells for 6 days at 30 °C. Conditioned media was dialysed against 2x phosphate buffered saline pH 7.4 buffer (Sigma-Aldrich). The spike ectodomain was purified using Strep-Tactin Superflow resin (IBA Lifesciences) followed by size exclusion chromatography using Superose 6 Increase column (GE Healthcare) equilibrated in 200 mM NaCl, 2 mM Tris-HCl pH 8.0, 0.02% NaN_3_ at 21 °C.

### SARS-CoV-2 S protein purification (Amsterdam)

The SARS-CoV-2 S protein construct was designed and expressed as described previously^1^. Briefly, a SARS-CoV-2 S gene encoding residues 1-1138 (WuhanHu-1; GenBank: MN908947.3) was ordered (Genscript) and cloned into a pPPI4 plasmid containing a T4 trimerization domain followed by a hexahistidine tag by PstI-BamHI digestion and ligation. Modifications consist of substituting the amino acid positions 986 and 987 to prolines and removing the furin cleavage site by substituting amino acids 682-685 to glycines. S protein was expressed in HEK 293F cells. Approximately 1.0 million cells/ml, maintained in Freestyle medium (Gibco), were transfected by adding a mix of PEImax (1 µg/µl) and SARS-CoV-2 S plasmid (312.5 µg/l) in a 3:1 ratio in OptiMEM. After six days supernatants were centrifuged for 30 min at 4000 rpm, and filtered using Steritop filter (Merck Millipore; 0.22 µm). S protein was purified from supernatant by affinity purification using Ni-NTA agarose beads and eluates were concentrated and buffer exchanged to PBS using Vivaspin filters with a 100 kDa molecular weight cutoff (GE Healthcare). The S protein concentration was determined by the Nanodrop method using the proteins peptidic molecular weight and extinction coefficient as determined by the online ExPASy software (ProtParam).

### SARS-CoV-2 Spike Trimer protein production and purification from CHO cells

For CHO-based production of a trimeric Spike protein the construct Spike_ΔCter_ΔFurin_2P_T4_His was used^53^. Briefly, based on the CHO-vector pXLG-6 (ExcellGene SA), containing a puromycin resistance marker and optimized expression elements, the variant spike sequence was inserted, containing a scambled furin cleavage site sequence, a two-prolin sequence for blocking the protein in the prefusion form, a 3’ T4-trimerization DNA, followed by a hexahistidin tag sequence. From chemical transfection (CHO4Tx®, ExcellGene) to clonal selection and isolation of protein producing cell lines, to subsequent expansion of cells and production – all steps were done under suspension culture in chemically defined media with strict adherence to regulatory recommendations.

Productions were done in orbitally shaken containers (TubeSpin bioreactor 600, TPP, Trasadingen, Switzerland) or in 5 L Erlenmeyer type shaker flasks in a Kuhner Shaker ISF4-X incubator (Adolf Kühner AG, Birsfelden, Switzerland), set to 37ºC, 5% CO2, humidity control and shaking at 150 rpm with a displacement radius of 50 mm. A fed-batch process was implemented until harvest of supernatants on day 10. The viability of cell culture, reaching about 25 × 106 cells/mL remained in the 90-100% range until harvest. Production culture fluids were subjected to purification by affinity chromatography after depth filtration to remove cells. Loading onto, washing of and elution from a Ni-Sepharose column (Cytiva) were optimized, following the resin producers’ suggestions. The eluted product stream was loaded on a Size-Exclusion column (SEC, Superdex 200 pg, Cytiva) for further purification, following a concentration step through tangential flow filtration.

### Plasmid construction and expression of Receptor Binding domain proteins

To express the receptor-binding domains (RBDs) of SARS-CoV-2 (residues: 320-527), SARS-CoV-1 (residues: 307-513), and MERS-CoV (residues: 368-587), we amplified the DNA fragments by PCR using the SARS-CoV-2, SARS-CoV-1 and MERS-CoV S protein ectodomain plasmids as templates. The DNA fragments were cloned into the phCMV3 (Genlantis, USA) vector in frame with the Tissue Plasminogen Activator (TPA) leader sequence. To facilitate the purification and biotinylation of the protein, we fused the protein with C-terminal 6X HisTag, and AviTag spaced by GS-linkers.

The RBD constructs were expressed by transient transfection of FreeStyle293F cells (Thermo Fisher). For a liter transfection, we added 350µg of the RBD-encoding plasmids to 16mL of Transfectagro (Corning) and 1.5mL of 40K PEI (1mg/mL) in 16mL of Transfectagro in separate 50mL conical tubes. The media containing the plasmids were filtered with a 0.22µm Steriflip Disposable Vaccum Filter Unit (Millipore Sigma) and gently mixed with PEI. The plasmid and PEI mixture was incubated for 30 minutes and gently added to 1L of cells at a concentration of 1million cells/mL. The culture supernatants were harvested after 4 days and loaded onto HisPur Ni-NTA resin column (Thermo Fisher). Following a wash with 25mM Imidazole, the protein from the column was eluted with 250mM Imidazole. The eluate was buffer exchanged with PBS and concentrated using 10K Amicon tuebs. The RBD proteins were further purified with size-exclusion chromatography using a Superdex 200 Increasse 10/300 GL column (GE Healthcare).

### Sample preparation and analysis by LC-MS

Mass spectrometry data for the virion derived glycosylation was obtained using the protocols described in Yao et al. and reanalyzed^33^. For recombinant S protein and RBD, 30 μg aliquots of SARS-CoV-2 S proteins were denatured for 1h in 50 mM Tris/HCl, pH 8.0 containing 6 M of urea and 5 mM of dithiothreitol (DTT). Next, the S proteins were reduced and alkylated by adding 20 mM IAA and incubated for 1hr in the dark, followed by incubation with DTT to get rid of any residual IAA. The alkylated S proteins were buffer-exchanged into 50 mM Tris/HCl, pH 8.0 using Vivaspin columns (3 kDa) and digested separately overnight using trypsin, chymotrypsin (Mass Spectrometry Grade, Promega) or alpha lytic protease (Sigma Aldrich) at a ratio of 1:30 (w/w). The peptides were dried and extracted using C18 Zip-tip (MerckMilipore). The peptides were dried again, re-suspended in 0.1% formic acid and analyzed by nanoLC-ESI MS with an Easy-nLC 1200 (Thermo Fisher Scientific) system coupled to an Orbitrap Fusion mass spectrometer (Thermo Fisher Scientific) using higher energy collision-induced dissociation (HCD) fragmentation. Peptides were separated using an EasySpray PepMap RSLC C18 column (75 µm × 75 cm). A trapping column (PepMap 100 C18 3 μm (particle size), 75 μm × 2cm) was used in line with the LC prior to separation with the analytical column. The LC conditions were as follows: 275 minute linear gradient consisting of 0-32% acetonitrile in 0.1% formic acid over 240 minutes followed by 35 minutes of 80% acetonitrile in 0.1% formic acid. The flow rate was set to 200 nL/min. The spray voltage was set to 2.7 kV and the temperature of the heated capillary was set to 40 °C. The ion transfer tube temperature was set to 275 °C. The scan range was 400−1600 *m/z*. The HCD collision energy was set to 50%. Precursor and fragment detection were performed using Orbitrap at a resolution MS1 = 100,000. MS2 = 30,000. The AGC target for MS1 = 4e^5^ and MS2 = 5e^4^ and injection time: MS1 = 50 ms MS2 = 54 ms.

### Data processing of LC-MS data

Glycopeptide fragmentation data were extracted from the raw file Byos (Version 3.5; Protein Metrics Inc.). The following parameters were used for data searches in Byonic: The precursor mass tolerance was set at 4 ppm and 10 ppm for fragments. Peptide modifications included in the search include: Cys carbamidomethyl, Met oxidation, Glu → pyroGlu, Gln → pyroGln and N deamidation. For each protease digest a separate search node was used with digestion parameters appropriate for each protease (Trypsin RK, Chymotrypsin YFW and ALP TASV) using a semi-specific search with 2 missed cleavages. A 1% false discovery rate (FDR) was applied. All three digests were combined into a single file for downstream analysis. All charge states for a single glycopeptide were summed. The glycopeptide fragmentation data were evaluated manually for each glycopeptide; the peptide was scored as true-positive when the correct b and y fragment ions were observed along with oxonium ions corresponding to the glycan identified. The protein metrics 309 mammalian N-glycan library was modified to include sulfated glycans and phosphorylated mannose species, although no phosphorylated mannose glycans were detected on any of the samples analysed. The relative amounts (determined by comparing the XIC of each glycopeptide, summing charge states) of each glycan at each site as well as the unoccupied proportion were determined by comparing the extracted chromatographic areas for different glycotypes with an identical peptide sequence. Glycans were categorized according to the composition detected. HexNAc(2), Hex(9−3) was classified as M9 to M3. Any of these compositions that were detected with a fucose are classified as FM. HexNAc(3)Hex(5−6)Neu5Ac)(0-4) was classified as Hybrid with HexNAc(3)Hex(5-6)Fuc(1)NeuAc(0-1) classified as Fhybrid. Complex-type glycans were classified according to the number of processed antenna and fucosylation. Complex glycans are categorized as HexNAc(3)(X), HexNAc(3)(F)(X), HexNAc(4)(X), HexNAc(4)(F)(X), HexNAc(5)(X), HexNAc(5)(F)(X), HexNAc(6+)(X) and HexNAc(6+)(F)(X). Core glycans are any glycan smaller than HexNAc(2)Hex(3) Glycans were further classified in Figures 2 and 3 according to their processing state. A glycan was classified as fucosylated or sialylated if it contained at least one fucose or sialic acid residue. Agalactosylated glycans consisted of compositions in the above categories for complex-type glycans which possessed only 3 hexose residues, or if the composition was a hybrid-type glycan then this was increased to 5 hexoses. If the complex-type glycan did not contain sialic acid but more than 3/5 hexoses then it was counted as galactosylated. Sulfated glycans were included in the above categories, so a glycan can be in both galactosylated and sulfated categories, for example.

### Integrative modelling and molecular dynamics simulation

The model of S protein was built using Modeller 9.21^75^ with three structural templates: i) the cryo-EM structure of SARS-CoV-2 S ECD in the open state (PDB: 6VSB)^29^, ii) the NMR structure of SAR-CoV S HR2 domain (PDB: 2FXP) ^61^, and iii) the NMR structure of HIV-1 gp-41 TM domain (PDB: 5JYN)^62^. Missing loops in the N-terminal domain and the C-terminal of the ECD were modelled using the cryo-EM structure of S ECD in the closed state, which was resolved at a higher resolution (PDB: 6XR8)^76^. A total of ten models were built and three top models (based on the lowest discrete optimized protein energy score^77^ were subjected to further stereochemical assessment using Ramachandran analysis^78^. The model with the lowest number of outliers was then selected. Man-9 glycans were added to all 22 glycosylation sites using the CHARMM-GUI Glycan Reader and Modeller web server^79^. Three cysteine residues on the C-terminal domain (C1236, C1240 and C1243) were palmitoylated as they have been shown to be important in cell fusion ^80^. The glycosylated S protein model was then embedded into a model ERGIC membrane built using CHARMM-GUI Membrane Builder^81^, which contained 47% phosphatidylcholine (PC), 20% phosphatidylethanolamine (PE), 11% phosphatidylinositol phosphate (PIP), 7% phosphatidylserine (PS) and 15% cholesterol^82–84^.

The system was parametrized using the CHARMM36 force field ^85^. TIP3P water molecules were used to solvate the system and 0.15 M NaCl salt was added to neutralize it. The system was then subjected to stepwise energy minimization and equilibration with decreasing positional and dihedral restraints, following the standard CHARMM-GUI protocol ^86^. A 200 ns production simulation was conducted with the temperature maintained at 310 K using the Nosé-Hoover thermostat ^87,88^. The pressure was maintained at 1 atm using a semi-isotropic coupling to the Parrinello-Rahman barostat ^89^. Electrostatics were calculated using the smooth particle mesh Ewald method ^90^ with a real-space cut-off of 1.2 nm, while the van der Waals interactions were truncated at 1.2 nm with a force switch smoothing function applied between 1.0-1.2 nm. The LINCS algorithm was utilized to constrain all covalent bonds involving hydrogen atoms ^91^ and a 2 fs integration time step was employed. All simulations were performed using GROMACS 2018 ^92^ and visualized in VMD ^93^. ASA and glycan contact analyses were performed using the built-in GROMACS tools *gmx sasa* and *gmx select*.

