## Supplementary Information for "Site-specific steric control of SARS-CoV-2 spike glycosylation"

[1] School of Biological Sciences, University of Southampton, Southampton, SO17 1BJ, UK
[2] Bioinformatics Institute, Agency for Science, Technology and Research (A*STAR), Singapore, 138671, Singapore
[3] Department of Immunology and Microbiology, The Scripps Research Institute, La Jolla, CA 13 92037, USA
[4] IAVI Neutralizing Antibody Center, The Scripps Research Institute, La Jolla, CA 92037, USA
[5] Consortium for HIV/AIDS Vaccine Development (CHAVD), The Scripps Research Institute, La Jolla, CA 92037, USA
[6] Department of Medical Microbiology, Amsterdam UMC, University of Amsterdam, Amsterdam Infection & Immunity Institute, Amsterdam, The Netherlands
[7] Tsinghua-Peking Center for Life Sciences, School of Life Sciences, Tsinghua University, Beijing 100084, China
[8] Beijing Advanced Innovation Center for Structural Biology & Frontier Research Center for Biological Structure, Beijing 100084, China
[9] Division of Molecular Medicine, Boston Children's Hospital, 3 Blackfan Street, Boston, MA 02115, USA
[10] Department of Pediatrics, Harvard Medical School, 3 Blackfan Street, Boston, MA 02115, USA
[11] Division of Structural Biology, University of Oxford, The Wellcome Centre for Human Genetics, Headington, Oxford, OX3 7BN, UK
[12] ExcellGene SA, CH1870 Monthey, Switzerland
[13] Biognos AB, Generatorsgatan 1 41705, Göteborg, Sweden
[14] Diamond Light Source Ltd, Harwell Science & Innovation Campus, Didcot, OX11 0DE, UK
[15] Department of Microbiology and Immunology, Weill Medical College of Cornell University, New York, NY, USA
[16] Ragon Institute of Massachusetts General Hospital, Massachusetts Institute of Technology, and Harvard University, Cambridge, MA 02139, USA
[17] Department of Biological Sciences, National University of Singapore, Singapore 117543, Singapore
[18] Department of Chemistry, Faculty of Science and Engineering, Manchester Institute of Biotechnology, The University of Manchester, Manchester M1 7DN, UK
* These authors contributed equally

†Correpsondence: Max Crispin and Peter Bond.

### Supplementary table 1: Average recombinant and viral-derived S protein site-specific glycan compositions averaged

| Averaged | N17 | N61 | N74 | N122 | N149 | N165 | N234 | N282 | N331 | N343 | N603 | N616 | N657 | N709 | N717 | N801 | N1074 | N1098 | N1134 | N1158 | N1173 | N1194 |
|---|---|---|---|---|---|---|---|---|---|---|---|---|---|---|---|---|---|---|---|---|---|---|
| M9Glc | 0 | 0 | 0 | 0 | 0 | 0 | 0 | 0 | 0 | 0 | 0 | 0 | 0 | 0 | 0 | 0 | 0 | 0 | 0 | 0 | 0 | 0 |
| M9 | 0 | 0 | 0 | 0 | 0 | 0 | 22 | 0 | 0 | 0 | 0 | 0 | 0 | 0 | 1 | 1 | 0 | 1 | 0 | 0 | 0 | 0 |
| M8 | 0 | 2 | 0 | 0 | 0 | 0 | 28 | 0 | 0 | 0 | 1 | 0 | 0 | 3 | 1 | 2 | 1 | 1 | 0 | 0 | 0 | 0 |
| M7 | 0 | 2 | 0 | 3 | 0 | 0 | 15 | 0 | 0 | 0 | 2 | 0 | 0 | 8 | 14 | 5 | 1 | 2 | 0 | 0 | 0 | 0 |
| M6 | 0 | 6 | 0 | 1 | 1 | 1 | 5 | 0 | 0 | 0 | 3 | 2 | 0 | 12 | 13 | 3 | 2 | 1 | 0 | 0 | 0 | 0 |
| M5 | 4 | 26 | 1 | 12 | 3 | 17 | 7 | 3 | 4 | 2 | 17 | 2 | 3 | 26 | 17 | 16 | 14 | 4 | 3 | 13 | 7 | 6 |
| M4 | 0 | 2 | 0 | 0 | 2 | 0 | 1 | 0 | 0 | 0 | 2 | 0 | 0 | 0 | 3 | 3 | 0 | 0 | 0 | 0 | 0 | 0 |
| M3 | 0 | 1 | 0 | 0 | 0 | 0 | 0 | 0 | 0 | 0 | 0 | 0 | 0 | 0 | 0 | 0 | 0 | 0 | 0 | 0 | 0 | 0 |
| FM | 0 | 0 | 1 | 0 | 0 | 0 | 0 | 0 | 0 | 0 | 0 | 0 | 0 | 0 | 0 | 0 | 0 | 0 | 0 | 0 | 0 | 0 |
| Hybrid | 1 | 11 | 0 | 13 | 0 | 2 | 4 | 1 | 1 | 0 | 7 | 4 | 0 | 0 | 22 | 17 | 5 | 8 | 1 | 0 | 0 | 0 |
| Fhybrid | 2 | 1 | 1 | 3 | 1 | 1 | 1 | 1 | 1 | 1 | 4 | 3 | 1 | 4 | 6 | 1 | 5 | 2 | 2 | 0 | 2 | 0 |
| HexNAc(3)(x) | 0 | 5 | 0 | 7 | 0 | 1 | 1 | 1 | 0 | 0 | 5 | 4 | 0 | 0 | 7 | 4 | 3 | 3 | 0 | 0 | 0 | 1 |
| HexNAc(3)(F)(x) | 4 | 0 | 4 | 4 | 4 | 0 | 0 | 1 | 2 | 2 | 2 | 4 | 1 | 4 | 3 | 1 | 6 | 2 | 2 | 0 | 4 | 7 |
| HexNAc(4)(x) | 0 | 18 | 0 | 25 | 2 | 5 | 7 | 4 | 4 | 1 | 1 | 8 | 0 | 0 | 3 | 14 | 3 | 24 | 0 | 0 | 0 | 0 |
| HexNAc(4)(F)(x) | 40 | 9 | 30 | 18 | 43 | 32 | 1 | 30 | 52 | 58 | 34 | 39 | 43 | 24 | 8 | 24 | 24 | 15 | 39 | 16 | 15 | 16 |
| HexNAc(5)(x) | 0 | 11 | 0 | 5 | 1 | 4 | 0 | 10 | 0 | 0 | 0 | 6 | 1 | 0 | 0 | 2 | 1 | 13 | 0 | 0 | 0 | 13 |
| HexNAc(5)(F)(x) | 37 | 2 | 44 | 6 | 36 | 33 | 0 | 38 | 31 | 34 | 22 | 21 | 33 | 16 | 2 | 6 | 24 | 12 | 20 | 23 | 9 | 3 |
| HexNAc(6+)(x) | 0 | 1 | 0 | 0 | 0 | 0 | 6 | 1 | 0 | 0 | 0 | 0 | 0 | 0 | 0 | 0 | 0 | 8 | 0 | 0 | 0 | 0 |
| HexNAc(6+)(F)(x) | 5 | 2 | 19 | 0 | 6 | 3 | 0 | 10 | 4 | 1 | 0 | 5 | 12 | 3 | 0 | 0 | 8 | 5 | 15 | 47 | 27 | 18 |
| Unoccupied | 6 | 0 | 0 | 1 | 1 | 0 | 1 | 0 | 0 | 0 | 0 | 0 | 6 | 0 | 0 | 0 | 2 | 0 | 15 | 0 | 35 | 36 |
| Core | 0 | 1 | 0 | 0 | 0 | 0 | 0 | 0 | 0 | 0 | 0 | 0 | 0 | 0 | 0 | 1 | 0 | 0 | 0 | 0 | 1 | 0 |
| Oligomannose | 5 | 39 | 2 | 17 | 6 | 19 | 80 | 4 | 4 | 2 | 25 | 4 | 3 | 49 | 48 | 30 | 19 | 7 | 5 | 13 | 7 | 7 |
| Hybrid | 3 | 11 | 1 | 16 | 1 | 2 | 4 | 1 | 2 | 1 | 11 | 8 | 1 | 4 | 28 | 18 | 10 | 10 | 3 | 0 | 2 | 0 |
| Complex | 86 | 49 | 97 | 66 | 92 | 79 | 15 | 95 | 94 | 97 | 64 | 88 | 89 | 47 | 24 | 50 | 69 | 82 | 77 | 87 | 55 | 57 |
| Unoccupied | 6 | 0 | 0 | 1 | 1 | 0 | 1 | 0 | 0 | 0 | 0 | 0 | 6 | 0 | 0 | 0 | 2 | 0 | 15 | 0 | 35 | 36 |
| Fucose | 87 | 14 | 98 | 32 | 90 | 69 | 3 | 79 | 91 | 96 | 63 | 73 | 89 | 50 | 19 | 32 | 67 | 36 | 78 | 86 | 57 | 43 |
| NeuAc | 26 | 7 | 43 | 21 | 35 | 18 | 0 | 15 | 31 | 37 | 4 | 9 | 33 | 6 | 6 | 28 | 13 | 46 | 18 | 41 | 14 | 32 |
| Sulfation | 0 | 0 | 8 | 0 | 1 | 0 | 0 | 0 | 0 | 0 | 0 | 0 | 0 | 0 | 0 | 0 | 0 | 0 | 0 | 0 | 0 | 0 |

| Viral derived | N17 | N61 | N74 | N122 | N149 | N165 | N234 | N282 | N331 | N343 | N603 | N616 | N657 | N709 | N717 | N801 | N1074 | N1098 | N1134 | N1158 | N1173 | N1194 |
|---|---|---|---|---|---|---|---|---|---|---|---|---|---|---|---|---|---|---|---|---|---|---|
| M9Glc | 0 | 0 | 0 | 0 | 0 | 0 | 0 | 0 | 0 | 0 | 0 | 0 | 0 | 0 | 0 | 0 | 0 | 0 | 0 | 0 | 0 | 0 |
| M9 | 0 | 0 | 0 | 0 | 0 | 0 | 28 | 0 | 0 | 0 | 0 | 0 | 0 | 0 | 0 | 0 | 0 | 0 | 0 | 0 | 0 | 0 |
| M8 | 0 | 0 | 0 | 0 | 0 | 0 | 53 | 0 | 0 | 0 | 4 | 0 | 0 | 0 | 0 | 0 | 5 | 1 | 0 | 0 | 0 | 0 |
| M7 | 0 | 11 | 0 | 0 | 0 | 0 | 15 | 0 | 1 | 0 | 9 | 0 | 0 | 0 | 22 | 12 | 10 | 1 | 0 | 0 | 0 | 0 |
| M6 | 0 | 37 | 0 | 4 | 0 | 0 | 4 | 0 | 0 | 0 | 12 | 0 | 0 | 44 | 20 | 31 | 20 | 1 | 0 | 0 | 0 | 0 |
| M5 | 0 | 12 | 0 | 0 | 0 | 0 | 0 | 0 | 0 | 0 | 6 | 0 | 0 | 16 | 13 | 0 | 0 | 1 | 0 | 0 | 0 | 0 |
| M4 | 0 | 3 | 0 | 0 | 0 | 0 | 0 | 0 | 0 | 0 | 0 | 0 | 0 | 0 | 0 | 0 | 0 | 0 | 0 | 0 | 0 | 0 |
| M3 | 0 | 0 | 0 | 0 | 0 | 0 | 0 | 0 | 0 | 0 | 0 | 0 | 0 | 0 | 0 | 0 | 0 | 0 | 0 | 0 | 0 | 0 |
| FM | 0 | 0 | 0 | 0 | 0 | 0 | 0 | 0 | 0 | 0 | 0 | 0 | 0 | 0 | 0 | 0 | 0 | 0 | 0 | 0 | 0 | 0 |
| Hybrid | 0 | 10 | 0 | 37 | 0 | 0 | 0 | 2 | 2 | 0 | 20 | 16 | 0 | 6 | 44 | 33 | 19 | 22 | 0 | 0 | 3 | 0 |
| Fhybrid | 0 | 0 | 0 | 8 | 0 | 0 | 0 | 0 | 0 | 13 | 2 | 0 | 0 | 0 | 0 | 0 | 0 | 2 | 14 | 0 | 0 | 0 |
| HexNAc(3)(x) | 0 | 0 | 0 | 7 | 0 | 0 | 0 | 2 | 1 | 0 | 3 | 0 | 0 | 0 | 0 | 0 | 0 | 3 | 0 | 0 | 0 | 0 |
| HexNAc(3)(F)(x) | 0 | 0 | 1 | 0 | 0 | 0 | 0 | 2 | 2 | 0 | 6 | 0 | 0 | 0 | 0 | 0 | 3 | 1 | 0 | 0 | 0 | 0 |
| HexNAc(4)(x) | 0 | 21 | 0 | 25 | 0 | 100 | 0 | 24 | 16 | 0 | 19 | 47 | 0 | 14 | 1 | 24 | 17 | 16 | 0 | 0 | 12 | 0 |
| HexNAc(4)(F)(x) | 34 | 0 | 35 | 17 | 57 | 0 | 0 | 42 | 47 | 5 | 19 | 36 | 0 | 4 | 0 | 0 | 20 | 2 | 86 | 0 | 22 | 0 |
| HexNAc(5)(x) | 8 | 0 | 0 | 2 | 0 | 0 | 0 | 5 | 7 | 11 | 0 | 0 | 0 | 0 | 0 | 0 | 6 | 38 | 0 | 0 | 18 | 0 |
| HexNAc(5)(F)(x) | 57 | 5 | 48 | 0 | 43 | 0 | 0 | 16 | 20 | 62 | 0 | 0 | 0 | 16 | 0 | 0 | 1 | 6 | 0 | 0 | 8 | 100 |
| HexNAc(6+)(x) | 0 | 0 | 0 | 0 | 0 | 0 | 0 | 0 | 0 | 9 | 0 | 0 | 0 | 0 | 0 | 0 | 0 | 7 | 0 | 0 | 0 | 0 |
| HexNAc(6+)(F)(x) | 0 | 0 | 16 | 0 | 0 | 0 | 0 | 7 | 2 | 0 | 0 | 0 | 0 | 0 | 0 | 0 | 0 | 0 | 0 | 0 | 38 | 0 |
| Unoccupied | 0 | 0 | 0 | 0 | 0 | 0 | 0 | 0 | 0 | 0 | 0 | 0 | 0 | 0 | 0 | 0 | 0 | 0 | 0 | 0 | 0 | 0 |
| Core | 0 | 0 | 0 | 0 | 0 | 0 | 0 | 0 | 1 | 0 | 0 | 0 | 0 | 0 | 0 | 0 | 0 | 0 | 0 | 0 | 0 | 0 |

| | | | | | | | | | | | | | | | | | | | | | | |
|---|---|---|---|---|---|---|---|---|---|---|---|---|---|---|---|---|---|---|---|---|---|---|
| Oligomannose | 0 | 63 | 0 | 4 | 0 | 0 | 100 | 0 | 1 | 0 | 31 | 0 | 0 | 60 | 55 | 44 | 35 | 4 | 0 | 0 | 0 | 0 |
| Hybrid | 0 | 10 | 0 | 45 | 0 | 0 | 0 | 3 | 2 | 13 | 22 | 16 | 0 | 6 | 44 | 33 | 19 | 24 | 14 | 0 | 3 | 0 |
| Complex | 100 | 27 | 100 | 51 | 100 | 100 | 0 | 97 | 96 | 87 | 47 | 84 | 0 | 34 | 1 | 24 | 47 | 73 | 86 | 0 | 97 | 100 |
| Unoccupied | 0 | 0 | 0 | 0 | 0 | 0 | 0 | 0 | 0 | 0 | 0 | 0 | 0 | 0 | 0 | 0 | 0 | 0 | 0 | 0 | 0 | 0 |
| Fucose | 92 | 5 | 100 | 25 | 100 | 0 | 0 | 67 | 71 | 80 | 27 | 36 | 0 | 20 | 0 | 0 | 24 | 10 | 100 | 0 | 68 | 100 |
| NeuAc | 0 | 0 | 0 | 0 | 0 | 0 | 0 | 0 | 2 | 0 | 0 | 0 | 0 | 0 | 0 | 0 | 0 | 1 | 0 | 0 | 0 | 0 |
| Sulfation | 0 | 0 | 0 | 0 | 0 | 0 | 0 | 0 | 0 | 0 | 0 | 0 | 0 | 0 | 0 | 0 | 0 | 0 | 0 | 0 | 0 | 0 |

**Supplementary table 2: Percentage point change in glycosylation on viral-derived S protein compared to recombinant S protein**

| | N17 | N61 | N74 | N122 | N149 | N165 | N234 | N282 | N331 | N343 | N603 | N616 | N657 | N709 | N717 | N801 | N1074 | N1098 | N1134 | N1158 | N1173 | N1194 | Average |
|---|---|---|---|---|---|---|---|---|---|---|---|---|---|---|---|---|---|---|---|---|---|---|---|
| M9Glc | 0 | 0 | 0 | 0 | 0 | 0 | 0 | 0 | 0 | 0 | 0 | 0 | | 0 | 0 | 0 | 0 | 0 | 0 | 0 | 0 | 0 | 0 |
| M9 | 0 | 0 | 0 | 0 | 0 | 0 | 5 | 0 | 0 | 0 | 0 | 0 | | 0 | -1 | -1 | 0 | -1 | 0 | 0 | 0 | 0 | 0 |
| M8 | 0 | -2 | 0 | 0 | 0 | 0 | 25 | 0 | 0 | 0 | 3 | 0 | | -3 | -1 | -2 | 4 | 0 | 0 | 0 | 0 | 0 | 1 |
| M7 | 0 | 10 | 0 | -3 | 0 | 0 | 0 | 0 | 0 | 0 | 6 | 0 | | -8 | 8 | 7 | 8 | 0 | 0 | 0 | 0 | 0 | 1 |
| M6 | 0 | 31 | 0 | 3 | -1 | -1 | -2 | 0 | 0 | 0 | 10 | -2 | | 32 | 7 | 29 | 18 | 0 | 0 | 0 | 0 | 0 | 6 |
| M5 | -4 | -14 | -1 | -12 | -3 | -17 | -7 | -3 | -4 | -2 | -11 | -2 | | -10 | -3 | -16 | -14 | -3 | -3 | -13 | -7 | -6 | -7 |
| M4 | 0 | 1 | 0 | 0 | -2 | 0 | -1 | 0 | 0 | 0 | -2 | 0 | | 0 | -3 | -3 | 0 | 0 | 0 | 0 | 0 | 0 | -1 |
| M3 | 0 | -1 | 0 | 0 | 0 | 0 | 0 | 0 | 0 | 0 | 0 | 0 | | 0 | 0 | 0 | 0 | 0 | 0 | 0 | 0 | 0 | 0 |
| FM | 0 | 0 | -1 | 0 | 0 | 0 | 0 | 0 | 0 | 0 | 0 | 0 | | 0 | 0 | 0 | 0 | 0 | 0 | 0 | 0 | 0 | 0 |
| Hybrid | -1 | 0 | 0 | 24 | 0 | -2 | -4 | 1 | 1 | 0 | 14 | 12 | | 6 | 22 | 16 | 14 | 14 | -1 | 0 | 3 | 0 | 6 |
| Fhybrid | -2 | -1 | -1 | 5 | -1 | -1 | -1 | 0 | -1 | 12 | -3 | -3 | | -4 | -6 | -1 | -5 | -1 | 12 | 0 | -2 | 0 | 0 |
| HexNAc(3)(x) | 0 | -5 | 0 | 0 | 0 | -1 | -1 | 1 | 1 | 0 | -2 | -4 | | 0 | -7 | -4 | -3 | 0 | 0 | 0 | 0 | -1 | -1 |
| HexNAc(3)(F)(x) | -4 | 0 | -3 | -4 | -4 | 0 | 0 | 1 | 0 | -2 | 3 | -4 | | -4 | -3 | -1 | -3 | -1 | -2 | 0 | -4 | -7 | -2 |
| HexNAc(4)(x) | 0 | 3 | 0 | 0 | -2 | 95 | -7 | 21 | 13 | -1 | 18 | 39 | | 14 | -2 | 9 | 14 | -8 | 0 | 0 | 12 | 0 | 10 |
| HexNAc(4)(F)(x) | -6 | -9 | 5 | -1 | 14 | -32 | -1 | 12 | -5 | -54 | -15 | -3 | | -19 | -8 | -24 | -4 | -13 | 47 | -16 | 7 | -16 | -7 |
| HexNAc(5)(x) | 8 | -11 | 0 | -3 | -1 | -4 | 0 | -5 | 7 | 11 | 0 | -6 | | 0 | 0 | -2 | 5 | 24 | 0 | 0 | 18 | -13 | 1 |
| HexNAc(5)(F)(x) | 20 | 3 | 3 | -6 | 8 | -33 | 0 | -22 | -11 | 29 | -22 | -21 | | 0 | -2 | -6 | -24 | -6 | -20 | -23 | -2 | 97 | -2 |
| HexNAc(6+)(x) | 0 | -1 | 0 | 0 | 0 | 0 | -6 | -1 | 0 | 9 | 0 | 0 | | 0 | 0 | 0 | 0 | -1 | 0 | 0 | 0 | 0 | 0 |
| HexNAc(6+)(F)(x) | -5 | -2 | -3 | 0 | -6 | -3 | 0 | -3 | -2 | -1 | 0 | -5 | | -3 | 0 | 0 | -8 | -5 | -15 | -47 | 11 | -18 | -5 |
| Unoccupied | -6 | 0 | 0 | -1 | -1 | 0 | -1 | 0 | 0 | 0 | 0 | 0 | | 0 | 0 | 0 | -2 | 0 | -15 | 0 | -35 | -36 | -5 |
| Core | 0 | -1 | 0 | 0 | 0 | 0 | 0 | 0 | 1 | 0 | 0 | 0 | | 0 | 0 | -1 | 0 | 0 | 0 | 0 | -1 | 0 | 0 |

| | N17 | N61 | N74 | N122 | N149 | N165 | N234 | N282 | N331 | N343 | N603 | N616 | N657 | N709 | N717 | N801 | N1074 | N1098 | N1134 | N1158 | N1173 | N1194 | Average |
|---|---|---|---|---|---|---|---|---|---|---|---|---|---|---|---|---|---|---|---|---|---|---|---|
| Oligomannose | -5 | 24 | -2 | -12 | -6 | -19 | 20 | -4 | -4 | -2 | 6 | -4 | | 11 | 7 | 13 | 16 | -3 | -5 | -13 | -7 | -7 | 0 |
| Hybrid | -3 | -1 | -1 | 28 | -1 | -2 | -4 | 1 | 0 | 12 | 11 | 9 | | 2 | 16 | 15 | 9 | 14 | 11 | 0 | 1 | 0 | 6 |
| Complex | 14 | -23 | 3 | -15 | 8 | 21 | -15 | 3 | 2 | -10 | -17 | -4 | | -13 | -23 | -27 | -22 | -10 | 9 | -87 | 41 | 43 | -6 |
| Unoccupied | -6 | 0 | 0 | -1 | -1 | 0 | -1 | 0 | 0 | 0 | 0 | 0 | | 0 | 0 | 0 | -2 | 0 | -15 | 0 | -35 | -36 | -5 |
| Fucose | 5 | -9 | 2 | -7 | 10 | -69 | -3 | -12 | -19 | -16 | -36 | -37 | | -30 | -19 | -32 | -43 | -26 | 22 | -86 | 11 | 57 | -16 |
| NeuAc | -26 | -7 | -43 | -21 | -35 | -18 | 0 | -15 | -30 | -37 | -4 | -9 | | -6 | -6 | -28 | -13 | -45 | -18 | -41 | -14 | -32 | -21 |
| Sulfation | 0 | 0 | -8 | 0 | -1 | 0 | 0 | 0 | 0 | 0 | 0 | 0 | | 0 | 0 | 0 | 0 | 0 | 0 | 0 | 0 | 0 | 0 |

**Supplementary table 3: Site-specific glycan compositions of the two RBD sites, N331 and N343**

**N331**

| | Viral derived S | Recombinant S | Monomeric RBD |
|---|---|---|---|
| M9Glc | 0% | 0% | 0% |
| M9 | 0% | 0% | 0% |
| M8 | 0% | 0% | 0% |
| M7 | 1% | 0% | 0% |
| M6 | 0% | 0% | 0% |
| M5 | 0% | 1% | 0% |
| M4 | 0% | 0% | 0% |
| M3 | 0% | 0% | 0% |
| FM | 0% | 0% | 0% |
| Hybrid | 2% | 0% | 0% |
| Fhybrid | 0% | 0% | 0% |
| HexNAc(3)(x) | 1% | 0% | 0% |
| HexNAc(3)(F)(x) | 2% | 3% | 0% |
| HexNAc(4)(x) | 16% | 0% | 1% |
| HexNAc(4)(F)(x) | 47% | 49% | 1% |
| HexNAc(5)(x) | 7% | 0% | 5% |
| HexNAc(5)(F)(x) | 20% | 40% | 10% |
| HexNAc(6+)(x) | 0% | 0% | 48% |
| HexNAc(6+)(F)(x) | 2% | 6% | 34% |
| Unoccupied | 0% | 0% | 0% |
| Core | 1% | 0% | 0% |

**N343**

| | Viral derived S | Recombinant S | Monomeric RBD |
|---|---|---|---|
| M9Glc | 0% | 0% | 0% |
| M9 | 0% | 0% | 0% |
| M8 | 0% | 0% | 0% |
| M7 | 0% | 0% | 0% |
| M6 | 0% | 0% | 0% |
| M5 | 0% | 0% | 0% |
| M4 | 0% | 0% | 0% |
| M3 | 0% | 0% | 0% |
| FM | 0% | 0% | 0% |
| Hybrid | 0% | 0% | 0% |
| Fhybrid | 13% | 0% | 0% |
| HexNAc(3)(x) | 0% | 0% | 0% |
| HexNAc(3)(F)(x) | 0% | 1% | 1% |
| HexNAc(4)(x) | 0% | 1% | 0% |
| HexNAc(4)(F)(x) | 5% | 61% | 28% |
| HexNAc(5)(x) | 11% | 0% | 0% |
| HexNAc(5)(F)(x) | 62% | 32% | 50% |
| HexNAc(6+)(x) | 9% | 1% | 0% |
| HexNAc(6+)(F)(x) | 0% | 3% | 20% |
| Unoccupied | 0% | 0% | 0% |
| Core | 0% | 0% | 0% |

**N331**

Viral derived S

| Sialylated | Galactosylated | Agalactosylated | Mannose | Unoccupied |
|---|---|---|---|---|
| 1.70% | 94.46% | 3.84% | 0.67% | 0.00% |

Recombinant S

| Sialylated | Galactosylated | Agalactosylated | Mannose | Unoccupied |
|---|---|---|---|---|
| 27.59% | 39.26% | 31.66% | 1.37% | 0.00% |

Monomeric RBD

| Sialylated | Galactosylated | Agalactosylated | Mannose | Unoccupied |
|---|---|---|---|---|
| 49.28% | 47.45% | 3.27% | 0.47% | 0.00% |

**N343**

Viral derived S

| Sialylated | Galactosylated | Agalactosylated | Mannose | Unoccupied |
|---|---|---|---|---|
| 0.00% | 100.00% | 0.00% | 0.00% | 0.00% |

Recombinant S

| Sialylated | Galactosylated | Agalactosylated | Mannose | Unoccupied |
|---|---|---|---|---|
| 59.09% | 31.61% | 9.31% | 0.00% | 0.00% |

Monomeric RBD

| Sialylated | Galactosylated | Agalactosylated | Mannose | Unoccupied |
|---|---|---|---|---|
| 29.77% | 44.02% | 26.21% | 0.30% | 0.00% |

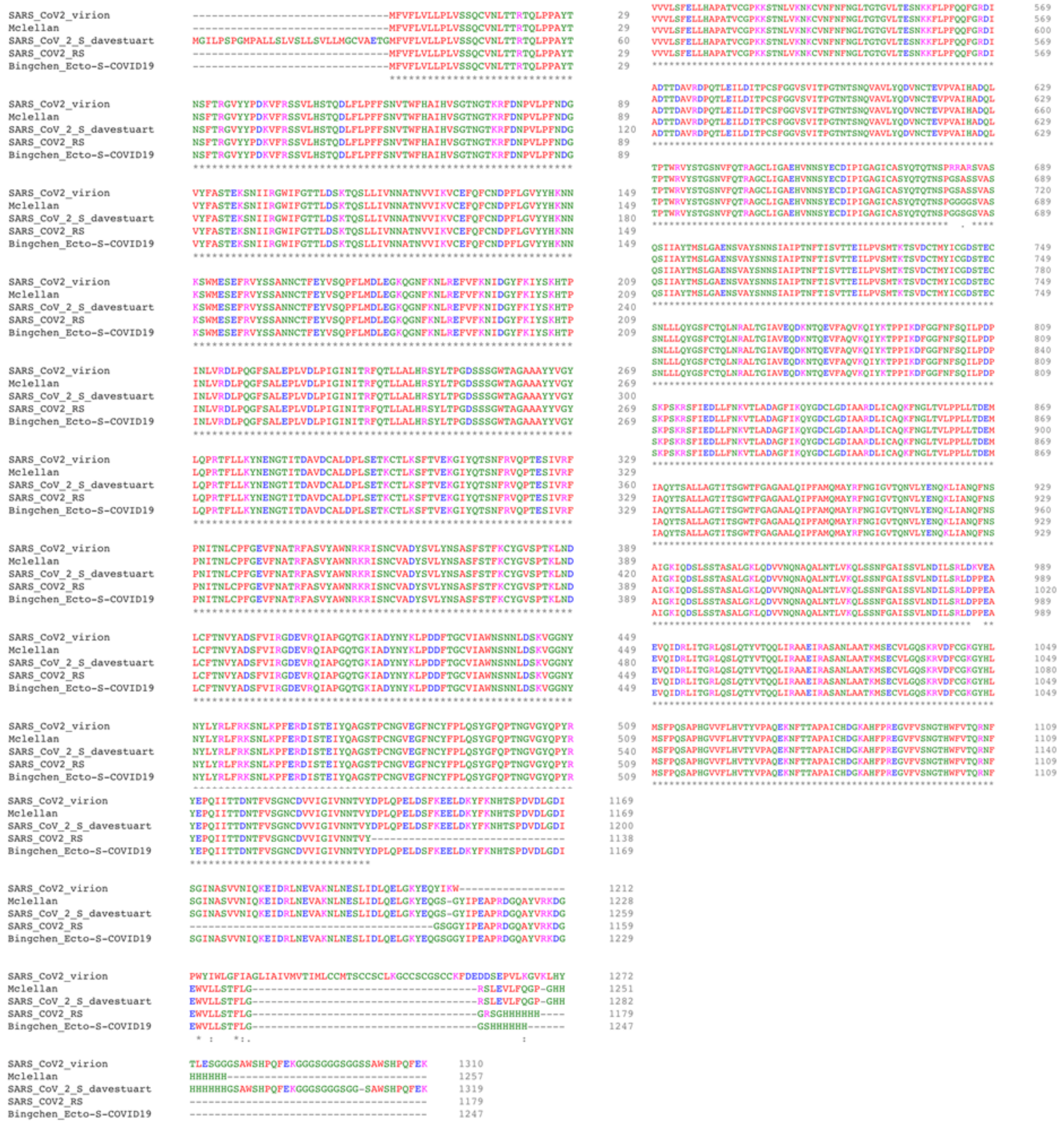

**Supplementary Figure 1: BLAST alignment of recombinant S proteins.** Sequences are labelled according to the principal investigator responsible for expression and purification. Protein analyzed from the binding site used the McLellan construct

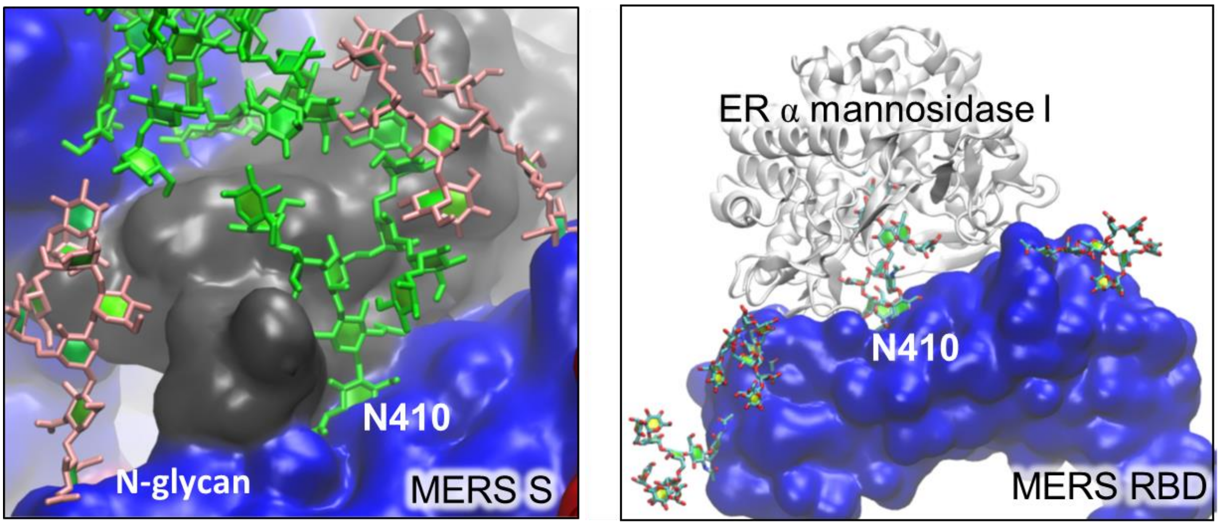

**Supplementary Figure 2: Model demonstrating the differential accessibility of the N410 glycan for glycan processing enzymes between trimeric MERS S protein and monomeric MERS RBD.**

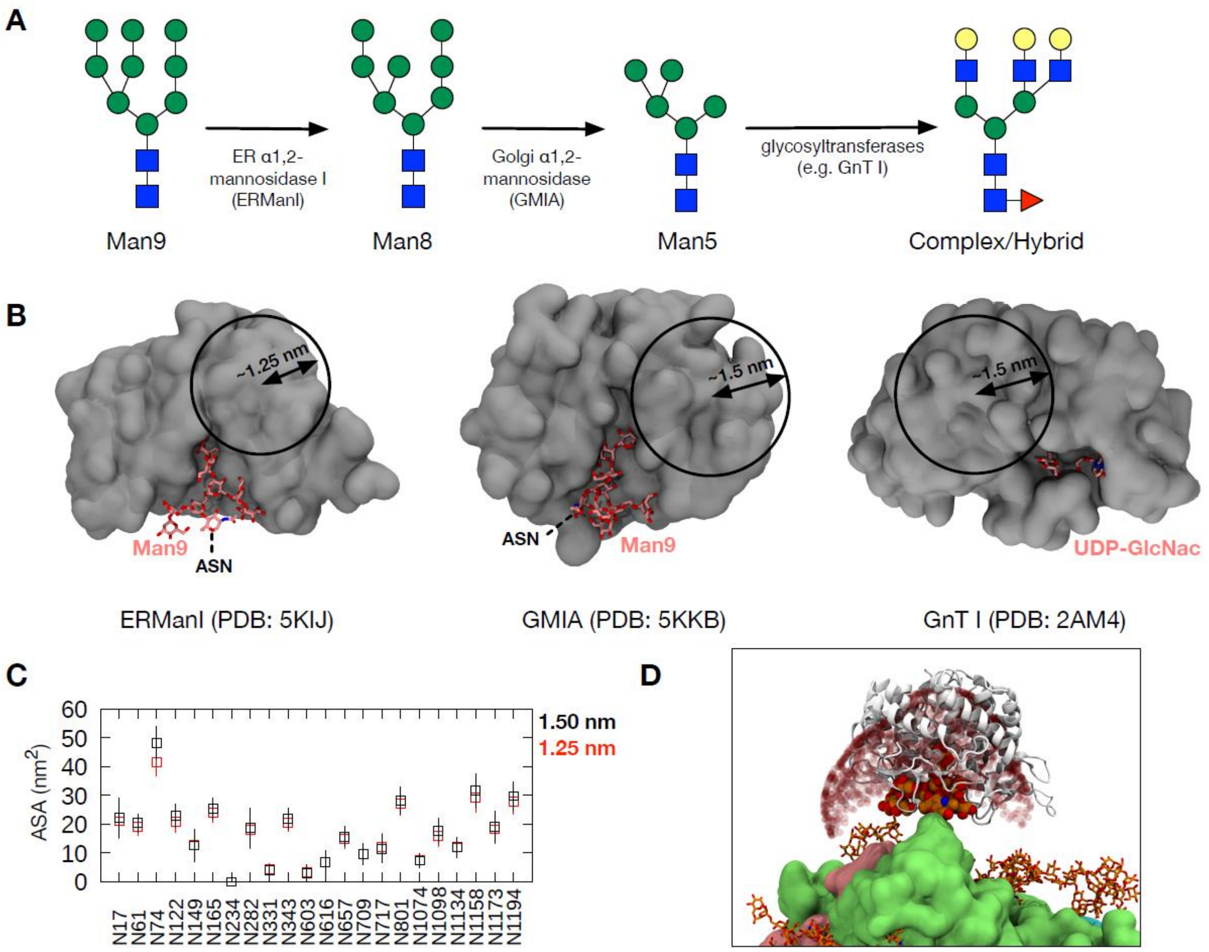

**Supplementary Figure 3: Validation of probe size for accessible surface area (ASA) measurement.** (A) A schematic of glycan processing from Man9 to complex/hybrid glycans and examples of enzymes involved. (B) Cross-section view of the crystal structures of these enzymes (grey in surface representation) bound to their substrate or substrate analogue (pink in stick representation). The circles show the approximate probe radii that could be used for glycan ASA calculation. (C) Comparison of average ASA values measured using 1.25 and 1.5 nm probe radii for Man9 in chain A of the S protein. Error bars indicate standard deviation along the simulation trajectory. (D) Accessible points (shown as red spheres) using a probe of 1.5 nm around glycan N74 (orange; van der Waals representation). S protein show in surface representation in green (chain A), cyan (chain B) and pink (chain C) and other glycans shown in stick representation in orange. ERManI enzyme shown in cartoon representation in white.

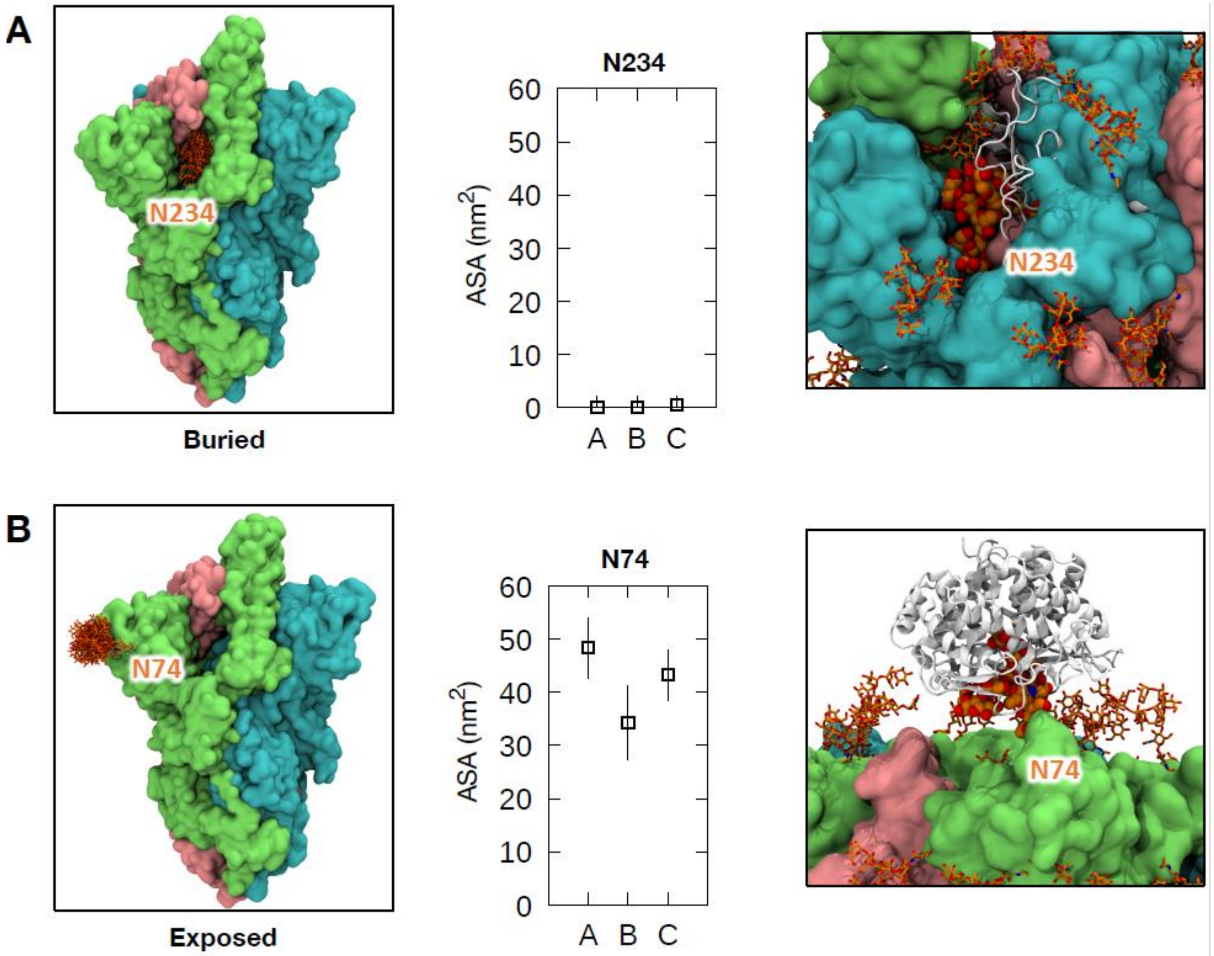

**Supplementary Figure 4: Examples of glycans either buried or exposed in all three chains.** (Left) Protein shown in surface representation with the individual chains coloured differently: chain A, green; chain B, cyan; chain C, pink. The overlay of glycan snapshots taken every 5 ns for the last 50 ns of the simulation is shown in stick representation. (Middle) Average ASA values for all three chains with error bars indicating the standard deviation along the trajectory. (Right) The structure of ERManI (PDB: 5KIJ) docked onto the glycans to show overlap with the S protein in A and full accessibility of the glycan in B. ERManI is shown in white and cartoon representation.

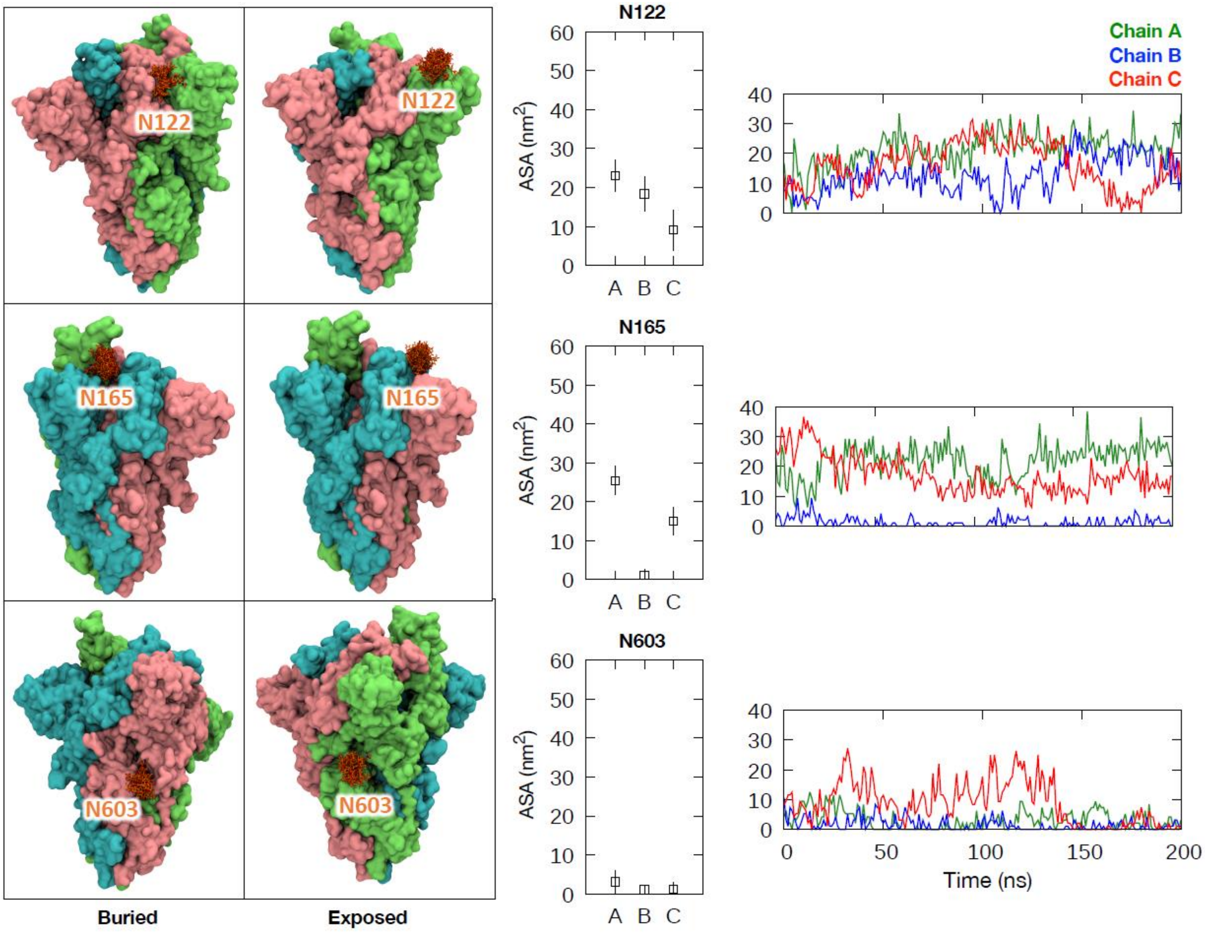

**Supplementary Figure 5: Examples of glycans with bimodal accessibility properties.** (Left) Protein is shown as in Supplementary Figure 3, with glycan snapshots taken from the portion of the simulation during which they were buried and exposed. (Middle) Average ASA values for all three chains with error bars indicating the standard deviation along the trajectory. (Right) ASA values for these glycans throughout the 200 ns simulation.

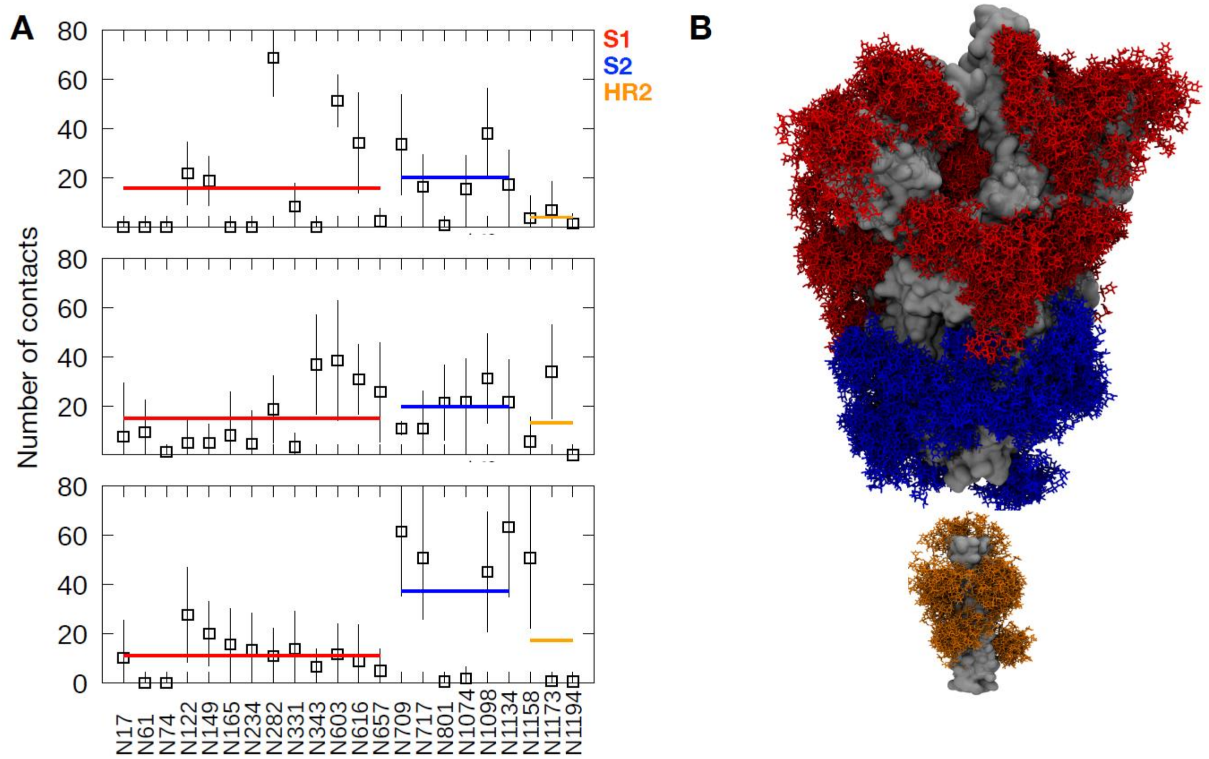

**Supplementary Figure 6: Glycan-glycan contacts from MD simulation.** (A) The average number of contacts made by glycans on each glycosylation site with any other glycans on the S protein throughout the last 50 ns of the simulation. The error bars show standard deviation along the trajectory. The thick red, blue and orange horizontal lines show the average contacts made by all glycans in the S1, S2 and HR2 regions, respectively. (B) An overlay of glycan snapshots taken every 10 ns along the 200 ns trajectory. Glycans are shown in stick representation and coloured red, blue and orange for S1, S2 and HR2 regions, respectively. S protein is shown in surface representation and coloured grey.
